# Adapting to heat stress by sowing summer grain crops early in late winter: Sorghum root growth, water use, and yield

**DOI:** 10.1101/2023.09.10.557017

**Authors:** Dongxue Zhao, Peter deVoil, Bethany G. Rognoni, Erin Wilkus, Joseph X Eyre, Ian Broad, Daniel Rodriguez

**Affiliations:** Centre for Crop Sciences, Queensland Alliance for Agriculture and Food Innovation (QAAFI), The University of Queensland, Gatton Campus, QLD 4343, Australia; Department of Agriculture and Fisheries (DAF), 13 Holberton Street, Toowoomba, QLD 4350, Australia; Department of Agriculture and Fisheries (DAF), 203 Tor Str Toowoomba, QLD 4350, Australia

**Keywords:** Climate adaption, Agronomy, Root morphology, Root phenotyping, Water use efficiency

## Abstract

**CONTEXT:** Drought and extreme heat at flowering are common stresses limiting the yield of summer crops, which are likely to intensify and become more frequent as projected under climate change.

**OBJECTIVE:** This study explores the idea that adaptation to these stresses could be increased by sowing summer crops early in late winter or spring, to avoid the overlap with critical crop stages around flowering. Here we report on the impacts of early sowing i.e., in late winter and spring on sorghum crop and root growth and function (i.e., water use), and final grain yield.

**METHODS:** Two seasons of on-farm genotype (G) by environment (E) by management (M) sorghum experimentation were conducted in the Darling Downs region of Queensland, Australia. Each trial consisted of a factorial combination of three times of sowing (TOS, referred to as late winter, spring, and summer), two levels of irrigation, four plant populations, and six commercial genotypes. Treatments were replicated three times. Crop roots and shoot were sampled at the flag leaf stage for each TOS. Crop water use across the growing season was monitored using time-lapse electromagnetic induction (EMI) surveys. EMI was also used to calculate a root activity factor. Final grain yield and yield components were determined at maturity.

**RESULTS:** Results showed that TOS, irrigation levels, and their interactions significantly influenced crop root and shoot traits, water use, and yield, though results were not always consistent across seasons. In the first season which was dry and had large temperature contrasts between TOS, crop growth in the early sown crops was primarily limited by temperature. In contrast, the second season was much warmer and crop growth was instead primarily limited by water availability. Cold air and soil temperatures in the early sowing dates i.e., late winter and spring during the first season, lead to smaller crops with smaller rooting systems and root-to-shoot ratios, and roots having a larger average root diameter. In general terms, root length and root length density responded positively to increasing pre-flowering mean air temperatures ranging between 16 and 20°C, while root average diameters were larger below 19 °C or above 21°C. Early sowing advanced flowering and therefore decreased the risk of extreme heat during the critical stages around flowering and affected water use before and after flowering. The root activity factor was directly related to the crop root length density. The early sown crops increased yield by transferring water use from vegetative to reproductive stages. The larger yield of the early sown crop was associated with larger grain numbers, particularly for the tillers, and a larger water use efficiency. As expected, irrigated and summer-sown crops exhibited lowest water use efficiency. The early-sown crops left more water in the soil profile at maturity, particularly under irrigated conditions and with small plant populations.

**CONCLUSIONS:** We conclude that early sown sorghum is a potential option to increase crop adaptation to hotter and drier environments. Here we propose that in the race to increase crop adaptation to heat stresses, plant breeding efforts should consider cold tolerance traits during crop germination, emergence, and early vegetative stages so that sorghum sowing windows could be significantly advanced.

## 1. Introduction

Sorghum (*sorghum bicolor* (L.) Moench) is a major dryland crop across Australia’s northern grains region, where droughts and extreme heat are common abiotic stresses limiting grain yield (Clarke et al., 2019). Across the region, and for conventional sorghum sowing times, there is a high frequency of heat affecting grain set (Singh et al., 2017). Even though heat stress affects multiple physiological processes i.e., photosynthesis, respiration, and transpiration (Prasad et al., 2017), the most yield sensitive phase to heat stress is concentrated around a narrow window i.e., 10–15 days around flowering (Singh et al., 2016). A short duration of high-temperature episodes coinciding with this window will damage an extensive series of morpho-physiological processes. In particular, pollen morphology damage (flattened and collapsed pollen) leads to reduced pollen viability and pollen germination on the stigmatic surface (Li et al., 2015). This causes failure of fertilization and seed set resulting in lower grain numbers and reduced grain yield (Singh et al., 2017). Terminal drought stresses after flowering likewise affect grain filling by reducing grain weight and quality (Prasad et al., 2015; Impa et al., 2019). Ongoing climate change is increasing global surface temperatures, the frequency and intensity of extreme heat and drought events (IPCC, 2021).

In-silico studies (Hammer et al., 2020) and empirical research (Borrell et al., 2000) showed that genotypes having reduced tillering, or agronomic practices that reduce water use early in the season, delay senescence and increase grain yield and yield quality (Thomas and Howarth, 2000). Pathways to increase adaptation to heat and drought stresses include improved genetic tolerance and agronomic avoidance (Prasad et al., 2015). Genetic tolerance to heat stress has been shown for both, the threshold at which pollen viability starts to be affected, and the response of pollen viability to increases in temperature above that threshold (Singh et al., 2015). In-silico assessments of the likely benefits of genetic tolerance to heat stress have shown yield gains between 5-8% and 13-17% under baseline and climate change projections, respectively (Singh et al., 2014). Clearly, in the long haul, plant breeding should be able to contribute to crop adaptation in warmer and drier environments (Nguyen et al., 2013).

Agronomy could mitigate heat stresses during flowering, and terminal drought stresses, by advancing flowering dates so that the overlap between times of the year of the high likelihood of the stresses and sensitive crop stages could be avoided. Sorghum crops sown in late winter or early spring will develop during periods of the year of lower atmospheric demand and flower before yield-limiting summer heat waves, reducing the impact of heat and terminal water stresses (Raymundo et al. 2021). However, sorghum is affected by cold temperatures early in the season (Marla et al., 2019). Sowing into soil temperatures lower than 16°C will slow the rate of metabolic activation enzymes in the seed (Patanè et al., 2021), leading to poor emergence, seedling establishment, and reduced plant stands (Rutayisire et al., 2021). Chilling temperatures after crop emergence will also reduce photosynthesis rates, and shoot and root growth. A poorly developed root system will likely limit access to soil water and nutrients (Aroca et al., 2001), consequently further constraining crop growth and production. Here we present results from a two-year on-farm research program in which we aimed to i) answer whether sowing sorghum into cold soils in late winter affects crop and root growth and function (i.e., water use), and final yield, and ii) study the relationships between ambient temperature, root traits, root function, shoot biomass, yield, and yield components. The study provides the first evidence relating root growth, root function, and yield across early- (i.e., late winter, spring) and summer-sown sorghum crops.

## 2. Materials and Methods

### 2.1. Study site and experimental design

On-farm trials were conducted at a commercial farm in Nangwee, QLD Australia (27°34’2.73" S, 151°18’34.36" E) during the 2019-2020 and 2020-2021 Southern Hemisphere summer growing seasons. The climate in the region is subtropical with an average of 621 mm rainfall per annum and mean annual maximum and minimum temperatures of 27.0 °C and 12.0 °C, respectively (Australian Government Bureau of Meteorology, 2021). The trial covered an area of ∼ 3.2ha (82m × 384m) of a uniform black, self-mulching cracking clay, characterized as a Vertosol soil (Isbell, 2016), with a clay content larger than 60%.

For each season, a split-plot experimental design was used for the random allocation of the factorial treatment structure of three times of sowing (TOS), two irrigation levels (rainfed/dryland and supplementary irrigation), six commercial genotypes, and four plant population densities (Table 1). There were three replicate blocks in each of the experiments. The factorial combinations of TOS and irrigation levels were randomly allocated to the mainplots, while the factorial combinations of genotypes and plant population densities were randomly allocated to subplots within the mainplots. The combination of treatments and replications generated 432 plots each 6m wide × 10m long. During the first season (2019-2020) the late winter sowing was on the 14^th^ of August, the spring sowing was on the 11^th^ of September, and the summer crop was sown on the 10^th^ of October. During the second season (2020-2021), the late winter, spring, and summer sorghum crops were sown on the 11^th^ of September, 6^th^ of October, and 5^th^ of November, respectively. The irrigated treatment was imposed by laying drip irrigation pipes along each row after sowing. Crops were fertilised following commercial sorghum production practices of the region and were kept free of weeds, pests and diseases. An automatic weather station and soil temperature probe were installed before sowing to monitor daily minimum and maximum temperature, soil temperature at seed depth, total radiation, and rainfall.

**Table 1.**
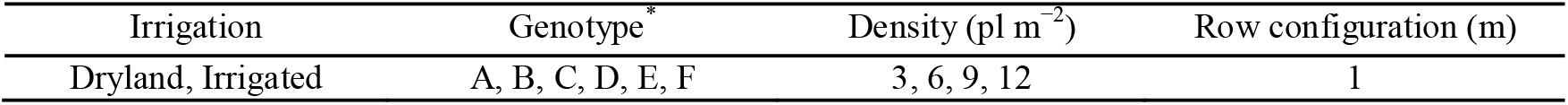
Description of treatments and crop configuration. Treatments include two levels of irrigation (rainfed/dryland and supplementary irrigation), six local commercial genotypes (coded), and four target plant population densities. The crop was sown in a 1 m row configuration.

| Irrigation | Genotype | Density (pl m<sup>-2</sup>) | Row configuration (m) |
| --- | --- | --- | --- |
| Dryland, Irrigated | A, B, C, D, E, F | 3, 6, 9, 12 | 1 |

### 2.2. Root and shoot growth

Roots and shoots from the TOS treatments were sampled at flag-leaf stage, on a single genotype (i.e., E, Table 1), a single plant population (9 pl m^-2^), and the two levels of irrigation, resulting in six plots being sampled per replication in each season. Shoots of twelve plants per plot were sampled and oven-dried at 75 °C until no further weight loss. After shoot sampling, the rooting system was sampled using a narrow tubular soil auger (44 mm diameter) down to a soil depth of 2.1 m. At each sampled plot, two soil cores (sub-samples) were taken in the interrow with each partitioned into eight depth intervals of 0-0.3, 0.3-0.5, 0.5-0.8, 0.8-1, 1-1.3, 1.3-1.5, 1.5-1.8 and 1.8-2.1m. The collected soil samples were soaked in water with a softening agent. The solution was then rinsed over a sieve in a root washing facility and the roots were collected with tweezers and stored in a 60-70% ethanol solution at 5°C. The root samples were then scanned using a digital scanner (Epson Expression XL 10000) with a resolution of 400 dpi. The scanned root images were analysed using the WinRHIZO^®^ software, Regent Instruments Inc., Quebec, Canada (Trachsel et al., 2011). Afterwards, the roots were dried at 75 °C for 72h and weighed to calculate root dry weight (g) at each depth.

The total root length (cm), average root diameter (cm), root surface area (cm^2^), and root volume (cm^3^) at each soil depth were calculated from WinRHIZO as in Rose (2017). The root length density (cm cm^-3^) and specific root length (cm g^-1^) at each depth were calculated by considering the sample soil volume and root dry weights.

The total root length (cm), total root surface area (cm^2^), total root dry weight (g), and total root volume (cm^3^) at plot level were then calculated by summing the corresponding root traits across the soil profile (0-2.1 m). The average root diameter at the plot level was determined from the total root length and total root volume. Similarly, at plot level average root length density (cm cm^-3^), average specific root length (cm g^-1^), and the root length to shoot dry weight ratio (cm g^-1^) were also calculated.

### 2.3. Dry matter production, yield, and yield components

Yield and biomass data were obtained on samples taken at grain physiological maturity from eight plants of central rows in each plot showing uniform plant density. Each sample was oven dried to a constant weight at 65 °C to determine the above-ground biomass. Panicles were then separated and threshed to determine yield components including grain number (grains m^-2^), grain weight (g per 1000 grains), and grain yield (t ha^-1^). Yield components were partitioned into main stems and tillers. The harvest index was estimated as the ratio of grain yield to biomass.

### 2.4. Measures of root function

Time-lapse electromagnetic induction (EMI) surveys were conducted to infer spatiotemporal variability of the soil volumetric water content (θ_v_, cm^3^ cm^-3^) and crop water use (mm) throughout the growing season. A DUALEM-21S (Dualem Inc., Milton, ON, Canada) instrument was used to collect soil apparent electrical conductivity (EC_a_), which is a function of soil moisture content. The instrument was towed 3m to the right of a four-wheel all-terrain vehicle that traversed the field along the transect in the middle of each plot. The vehicle was driven at a speed of ∼4 km h^-1^, thus approximately five EC_a_ reading were recorded per plot. In the first season, a total of five EMI surveys were collected (Fig. S1a). In the second season, a total of 17 EMI surveys were taken at weekly or fortnightly intervals (Fig. S1b).

A detailed description of the method used to convert EC_a_ to θ_v_ is in Zhao et al. (2022). The crop water use down to 1.5m was determined for every two consecutive EMI surveys (Δ*S*, mm) as in eq. 1:

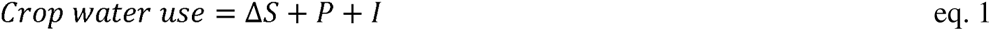

Where Δ*S* (mm) is the change in soil water content in the 0-1.5m, P is precipitation (mm) and I irrigation (mm). Crop water use was also divided into pre-flowering, post-flowering, and total crop water use. Water use efficiency (WUE, t ha^-1^ mm^-1^) was calculated as the ratio between grain yield (t ha^-1^) and total crop water use (mm).

In addition, a root activity factor at around flowering in the 2020-2021 was calculated to represent the presence and activity in each studied soil depth as in Zhao et al., 2022 (eq. 2) given the ample EMI surveys in this season. Briefly, eq. 2 assumes that water use from an *i^th^*soil layer can be represented by the plant available water (mm) of that *i^th^*soil layer, a term representing the size of the canopy, and a factor we call *root activity factor* (R) (eq. 2). Another assumption is that given the large volume of soil surveyed, that all treatments were affected by the same environmental conditions, and as all plots are measured within a small-time window (∼2hs), changes in atmospheric demand can be expected to be small.

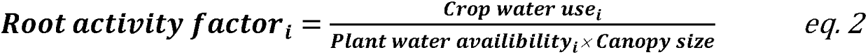

In eq. 2, *Root activity factor_i_* indicates the root presence and activity in the *ith* layer; *Water use_i_* is the change in water content (mm) in the *ith* soil layer between two consecutive EMI determinations around flowering and permanent crop wilting point; *Plant water availability_i_* is the plant available water (mm) in the *ith* layer at the start of the measurement period; and canopy size as main determinant of crop water demand. The Normalized Difference Vegetation Index (NDVI) was used as a proxy to account for canopy size. In this study, NDVI around flowering for each TOS was derived from satellite images from PlanetScope. For more details refer to Zhao et al., 2022.

### 2.5. Statistical analysis

Root traits were analysed using a linear mixed model (LMM) framework for each season at both plot and across depths. At the plot level, the LMMs included fixed effects for TOS, irrigation, and the interaction between TOS and irrigation. Design effects were included as random effects and consisted of a core nested structure of replicate, plots, and sub-samples.

Across depths, the LMMs were used to test the effects of TOS, irrigation, depth, and their interactions on root traits. The residual variance model was upgraded in stages, to test for heterogeneity of residual variance between depth intervals, as well as residual correlation models across depth intervals. The most parsimonious model for each measure was selected using the Akaike Information Criterion (Akaike, 1998). Moreover, the values of root traits (i.e., root length (cm), root surface area (cm^2^), root dry weight (g) and root volume (cm^3^)) were weighted on a “per 10 cm” basis to account for the differing widths of the depth intervals.

Effects of season, TOS, irrigation, plant population, genotype, and their interactions on the yield components (i.e., grain yield, grain number and grain weight) were also tested with LMMs. The season, TOS, irrigation, plant population, and genotype levels were used as fixed factors. Design effects were included as random effects. Separate residual variances were fitted for the two seasons. The predicted means for differences between the five-way interactions (season×TOS×irrigation×plant population×genotype) were derived for the yield components of the plant, main stem, and tiller, respectively. Then, the relationships between grain yield and grain number were analysed for each season independently using simple linear regression embedded in JMP 16 (SAS Institute, 2021). The 95% prediction interval was calculated to assess the prediction uncertainty of the linear regression.

All LMMs were fitted using the ‘ASReml-R’ statistical package (Butler et al. 2017), whereby variance components were estimated using residual maximum likelihood (Patterson & Thompson 1971) in R (R Core Team 2022). The fixed effects were tested using Wald tests (Kenward & Roger 1997), and Empirical Best Linear Unbiased Estimates (eBLUEs) were generated from the models for significant effects. Significant differences between pairs of treatments were determined using Fisher’s least significant difference (LSD) (Welham et al. 2014), and all significances were assessed at the 5% level.

To explore the environmental effects of TOS on root growth, root function (i.e., crop water use), yield components and harvest index, a principal component analysis (PCA) was performed including environmental covariates (Table 2), using the ‘stats’ package in R. In addition, the relationships between plot level root traits and pre-flowering mean air temperature and between WUE and yield components were also fitted in JMP 16 based on the least squares function.

**Table 2.**
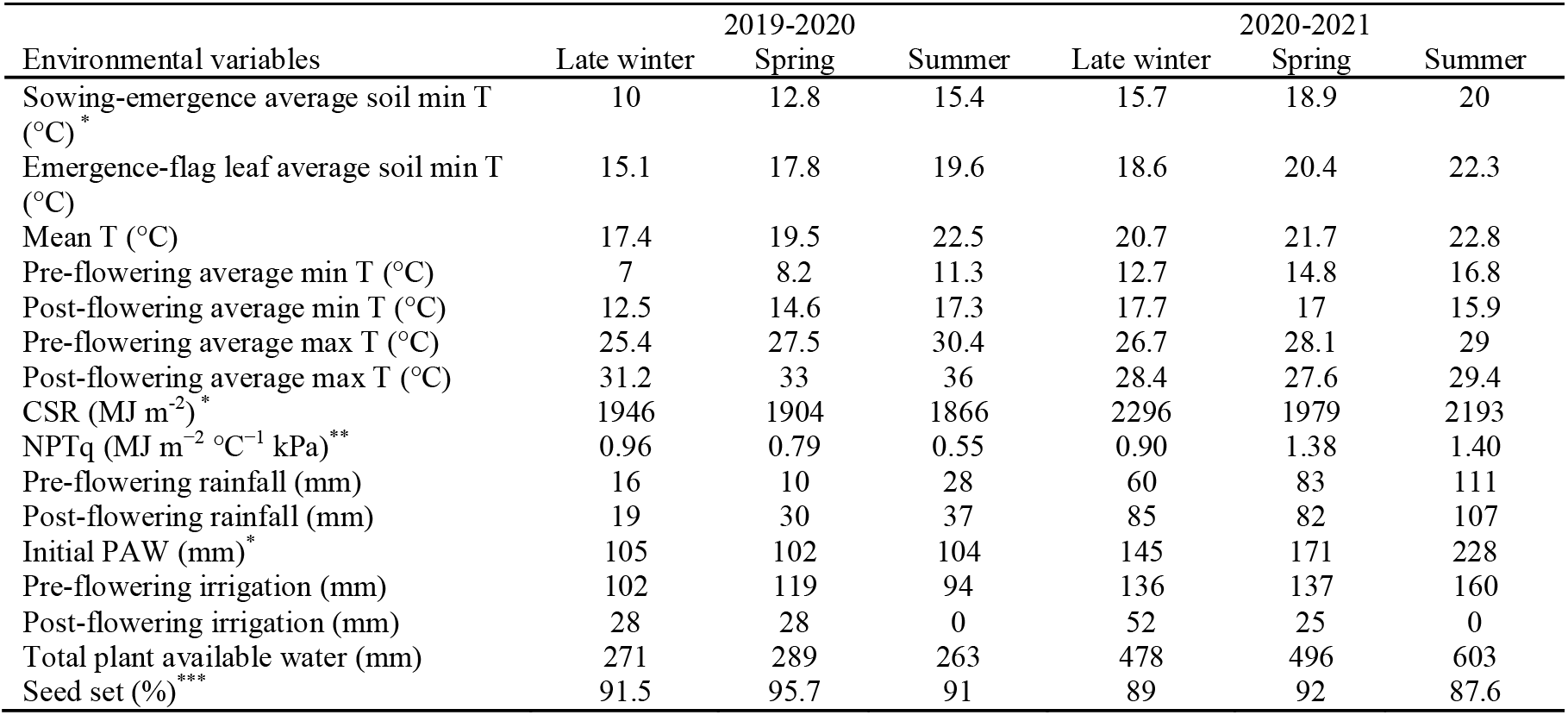
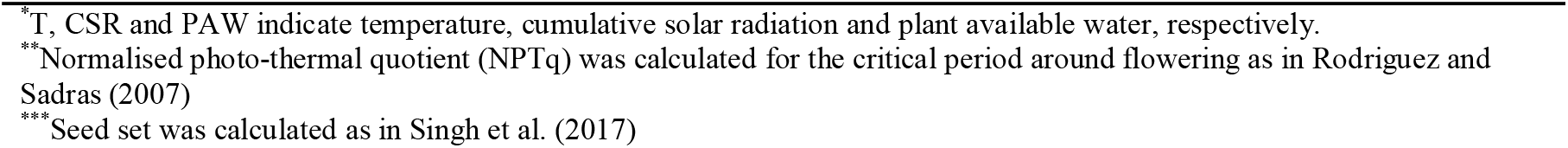
Environmental conditions for the late winter, spring, and summer sown sorghum in the 2019-2020 and 2020-2021 growing seasons at Nangwee, Queensland, Australia.

| Environmental variables | 2019-2020 |  |  | 2020-2021 |  |  |
| --- | --- | --- | --- | --- | --- | --- |
|  | Late winter | Spring | Summer | Late winter | Spring | Summer |
| Sowing-emergence average soil min T (°C)* | 10 | 12.8 | 15.4 | 15.7 | 18.9 | 20 |
| Emergence-flag leaf average soil min T (°C) | 15.1 | 17.8 | 19.6 | 18.6 | 20.4 | 22.3 |
| Mean T (°C) | 17.4 | 19.5 | 22.5 | 20.7 | 21.7 | 22.8 |
| Pre-flowering average min T (°C) | 7 | 8.2 | 11.3 | 12.7 | 14.8 | 16.8 |
| Post-flowering average min T (°C) | 12.5 | 14.6 | 17.3 | 17.7 | 17 | 15.9 |
| Pre-flowering average max T (°C) | 25.4 | 27.5 | 30.4 | 26.7 | 28.1 | 29 |
| Post-flowering average max T (°C) | 31.2 | 33 | 36 | 28.4 | 27.6 | 29.4 |
| CSR (MJ m<sup>-2</sup>)* | 1946 | 1904 | 1866 | 2296 | 1979 | 2193 |
| NPTq (MJ m<sup>-2</sup> °C<sup>-1</sup> kPa)** | 0.96 | 0.79 | 0.55 | 0.90 | 1.38 | 1.40 |
| Pre-flowering rainfall (mm) | 16 | 10 | 28 | 60 | 83 | 111 |
| Post-flowering rainfall (mm) | 19 | 30 | 37 | 85 | 82 | 107 |
| Initial PAW (mm)* | 105 | 102 | 104 | 145 | 171 | 228 |
| Pre-flowering irrigation (mm) | 102 | 119 | 94 | 136 | 137 | 160 |
| Post-flowering irrigation (mm) | 28 | 28 | 0 | 52 | 25 | 0 |
| Total plant available water (mm) | 271 | 289 | 263 | 478 | 496 | 603 |
| Seed set (%)*** | 91.5 | 95.7 | 91 | 89 | 92 | 87.6 |

\*T, CSR and PAW indicate temperature, cumulative solar radiation and plant available water, respectively.
\*\*Normalised photo-thermal quotient (NPTq) was calculated for the critical period around flowering as in Rodriguez and Sadras (2007)
\*\*\*Seed set was calculated as in Singh et al. (2017)

## 3. Results

### 3.1. Environmental conditions and crop phenology

The combination of season, TOS and irrigation exposed the crop to a highly diverse range of growing conditions (Table 2). In both seasons, soil temperatures for the late winter sown crop were below or close to the recommended 16°C at sowing depth. Minimum temperatures were particularly low during the first season, which was also drier than the second season. Moreover, the early sown crops were exposed to chilling ambient temperatures between emergence to flowering.

Growing conditions affected phenology when expressed in calendar days, though the differences were small when expressed in cumulative thermal units (Table 3). Early sown crops experienced a longer period between sowing and emergence in both seasons (Table 3). Table 3 shows that for the late winter sown sorghum, the period to emergence was extended by a factor of three. The lower temperatures for late winter sowing also contributed to a slightly longer period between emergence and flowering, indicating that the main delay in crop phenology for early sowing was driven by longer sowing to emergence and emergence to flowering periods. In comparison, the days from flowering to maturity were similar between different times of sowing in both seasons. Larger values of cumulative degree days were observed for the summer-sown sorghum between emergence and flowering, and emergence to maturity. There were no significant differences in phenology across genotypes in either season (Table S1).

**Table 3.** Sorghum phenology for late winter, spring, and summer sown crops during the 2019-2020 and 2020-2021 growing seasons at Nangwee, Queensland, Australia. Values indicate days and the ranges indicate variability between treatments other than time of sowing (TOS).

| Stages | TOS | 2019-2020 |  | 2020-2021 |  |
| --- | --- | --- | --- | --- | --- |
|  |  | Days | CGD (°C d) | Days | CGD (°C d) |
| Sowing-Emergence | Late winter | 18-28 | 73-120 | 8-11 | 52-83 |
|  | Spring | 14-17 | 119-152 | 5-6 | 56-67 |
|  | Summer | 6 | 77 | 5 | 46 |
| Emergence-Flowering | Late winter | 67-79 | 761-782 | 67-76 | 651-787 |
|  | Spring | 60-69 | 765-858 | 66-70 | 770-832 |
|  | Summer | 63-69 | 893-1004 | 67-71 | 845-903 |
| Emergence-Maturity | Late winter | 98-108 | 1086-1293 | 106-109 | 1153-1184 |
|  | Spring | 92-97 | 1256-1327 | 92-96 | 1066-1119 |
|  | Summer | 92-96 | 1363-1452 | 101-103 | 1302-1359 |

CGD is the cumulative growing degree days from sowing calculated using a base temperature of 10 °C and daily minimum and maximum temperatures from the automatic weather station installed in the field.

### 3.2. Roots and shoots growth

During the first season, the late winter sown sorghum had a significantly smaller total root length than the spring and summer sown sorghum crops (Fig. 1a). This was also the case for total root surface area (Fig. 1b) and average root length density (Fig. 1c). Conversely, the late winter sown crop had significantly thicker roots i.e., average root diameter than the spring and summer sown crops (Fig. 1d). Compared to spring and summer sown crops, late winter crops were smaller (Fig. 1f), particularly in the dryland (by 49-52%). Similarly, late winter crops had a smaller total root length to shoot dry weight ratio (Fig. 1e). In the second, warmer season (and later sown late winter crop), the value of the root traits was generally larger than in the first season, although there were no significant differences between treatments (Table 4).

**Figure 1.**
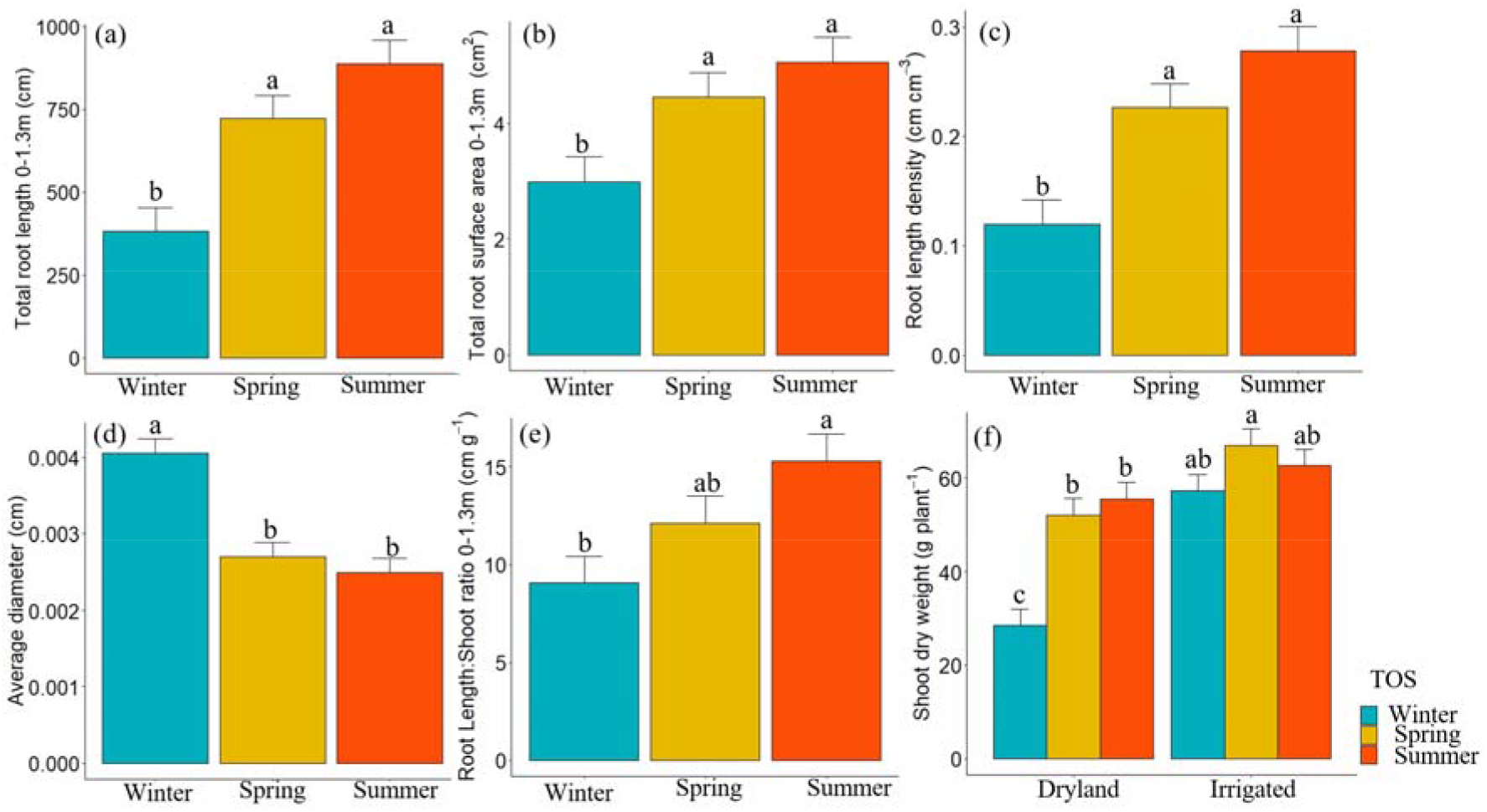
Effect of time of sowing (TOS) including late winter, spring, and summer on (a) total root length (cm), (b) total root surface area (cm^3^), (c) root length density (cm cm^-3^), (d) average root diameter (cm), (e) total root length to shoot weight (cm g^-1^) ratio and the effects of TOS by irrigation on (f) shoot dry weight at plot level at the flag leaf stage, for a sorghum crop (Genotype E) sown at 9 pl m^-2^ in the 2019-2020 season at Nangwee, Queensland, Australia. Different lowercase letters indicate a significant difference at *p* ≤ 0.05. Error bars represent standard errors of the estimations.

**Table 4.** Results from the linear mixed model of root traits, exploring the factorial combination of time of sowing (TOS, late winter, spring, and summer), and irrigation (dryland and irrigation), and their interactions for a sorghum crop (Genotype E) sown at 9 pl m^-2^ in the 2019-2020 and 2020-2021 seasons, respectively at Nangwee, Queensland, Australia.

|  | 2019-2020 |  |  |  | 2020-2021 |  |  |  |
| --- | --- | --- | --- | --- | --- | --- | --- | --- |
|  | Mean | Significance | df | p | Mean | Significance | df | p |
| Total root length (cm) | 663.1 | TOS main effect | 2 | 0.000*** | 927.7 | ns |  | > 0.05 |
| Average root diameter (mm) | 0.003 | TOS main effect | 2 | 0.000*** | 0.003 | ns |  | > 0.05 |
| Total root surface area (cm<sup>2</sup>) | 4.2 | TOS main effect | 2 | 0.014* | 5.6 | ns |  | > 0.05 |
| Average root length density (cm cm<sup>-3</sup>) | 0.208 | TOS main effect | 2 | 0.000*** | 0.291 | ns |  | > 0.05 |
| Total root length: shoot dry weight ratio (cm g<sup>-1</sup>) | 12.3 | TOS main effect | 2 | 0.023* | 16 | ns |  | > 0.05 |
| Shoot dry weight (g pl<sup>-1</sup>) | 54 | TOS × Irrigation main effect | 2 | 0.030* | 62 | ns |  | > 0.05 |
| Total root dry weight (g) | 0.135 | ns |  | > 0.05 | 0.145 | ns |  | > 0.05 |
| Total root volume (cm<sup>3</sup>) | 0.004 | ns |  | > 0.05 | 0.007 | ns |  | > 0.05 |
| Average specific root length (cm g<sup>-1</sup>) | 5100.7 | ns |  | > 0.05 | 6923.6 | ns |  | > 0.05 |

Where \*, \*\* and \*\*\* represent the significant level $\leq 0.05$ , $\leq 0.01$ and $\leq 0.001$ respectively, and ns indicates not significant.

Fig. 2 and Fig. 3 show root traits (eBLUEs) from the LMM for the Depth×TOS or Depth×TOS×Irrigation interactions. In the 2019-2020 season, late winter sowing significantly decreased root length (Fig. 2a) and root length density (Fig. 2b) at each soil depth. Whereas the opposite was true for the average diameter (Fig. 2c) in which late winter sown sorghum significantly increased the average root diameter at the 0-0.8 m soil profile. This was also the case for the root volume (Fig. 2d), especially in the dryland treatments. In contrast, late winter sowing reduced the surface area (Fig. 3a), root dry weight (Fig. 3b), and specific root length (Fig. 3c) across the soil profiles although the difference in these three traits between TOS was not significant.

**Fig. 2.**
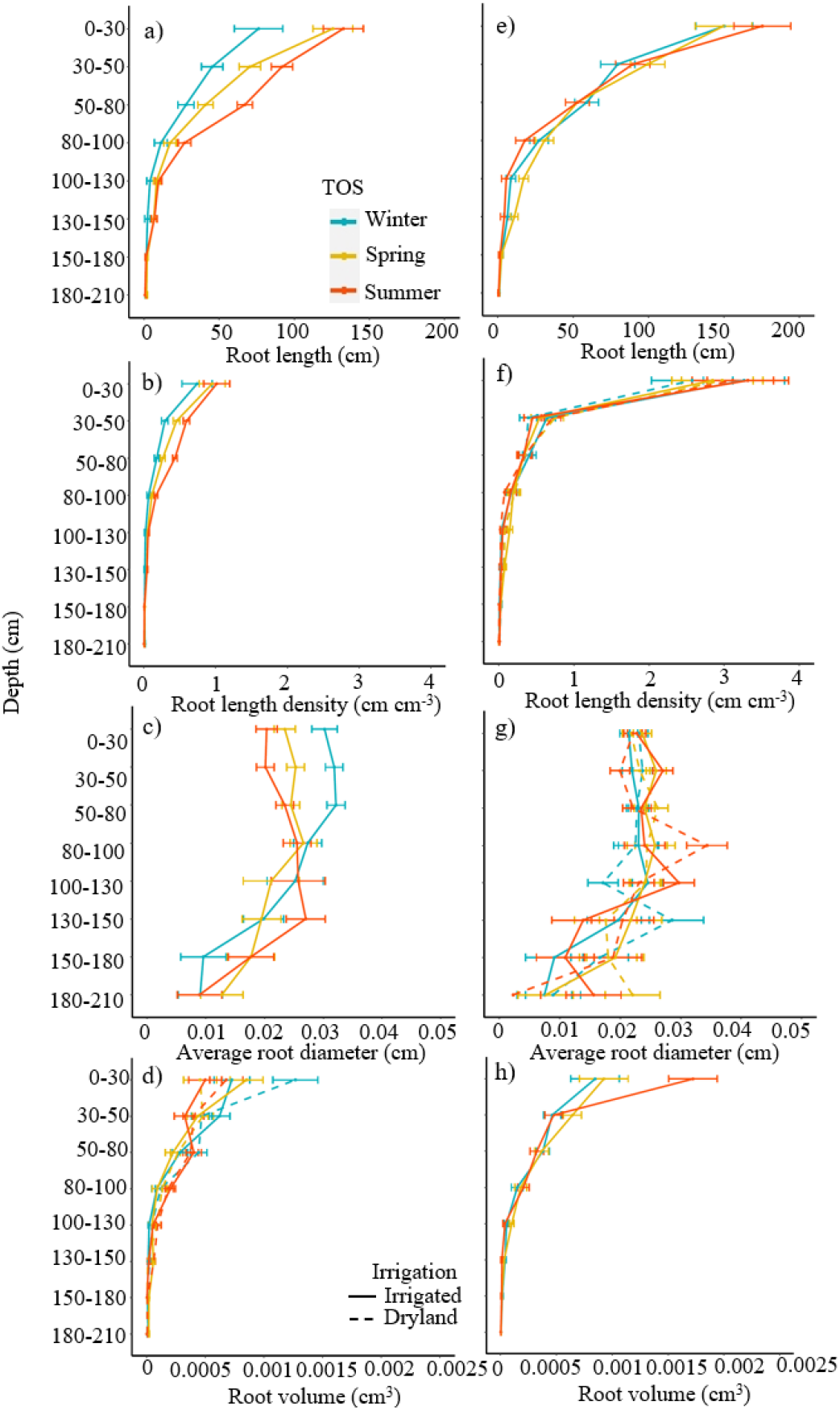
Effects of depth by the time of sowing (TOS including late winter, spring, and summer) or depth by TOS by irrigation on root length, root length density, average root diameter, and root volume in 2019-2020 (a, b, c, and d, respectively) and 2020-2021 seasons (e, f, g and h, respectively) at the flag leaf stage, for a sorghum crop (Genotype E) sown at 9 pl m^-2^ at Nangwee, Queensland, Australia. Values were the means for the three replicates. Error bars represent standard errors of the estimations.

**Figure 3.**
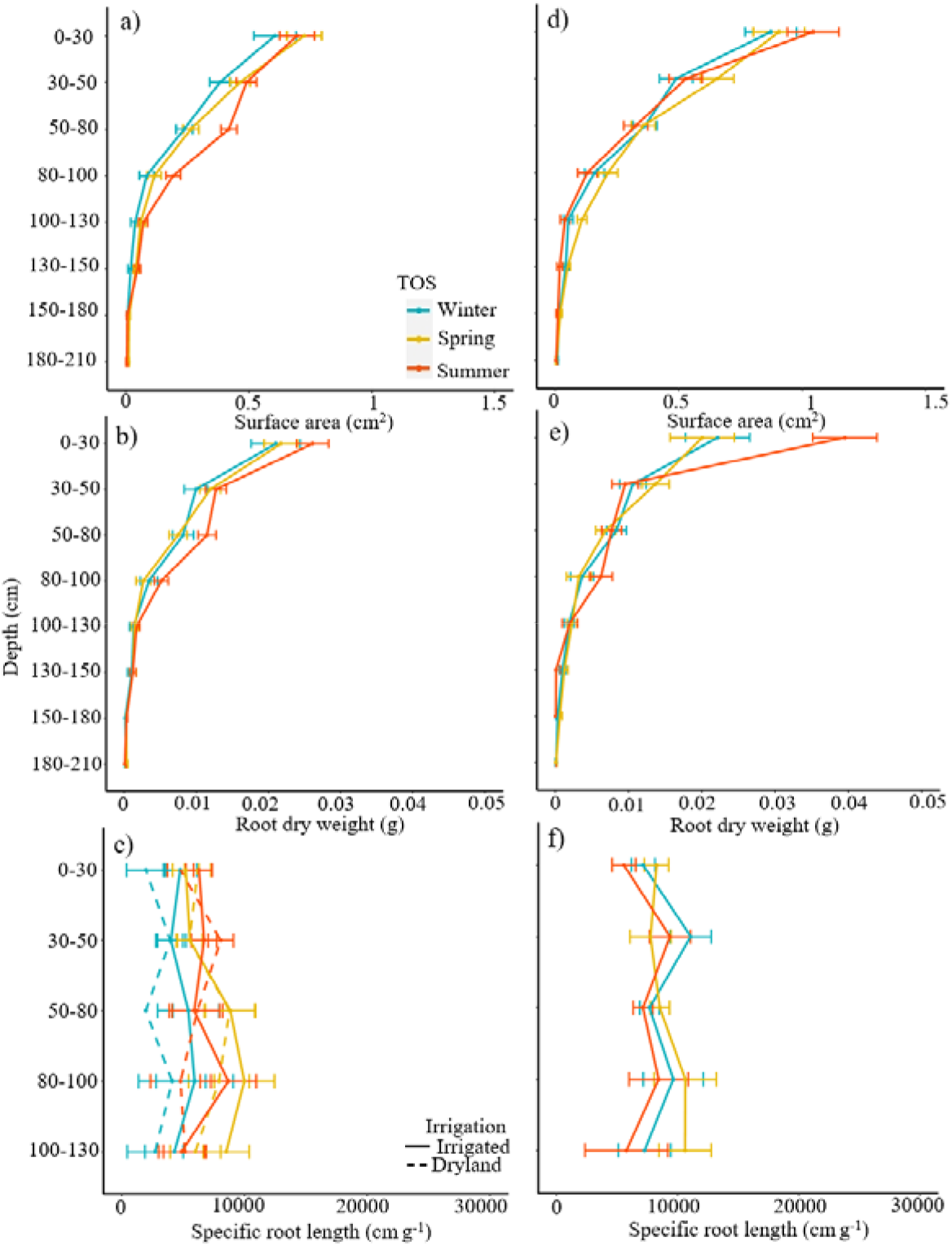
Effects of depth by the time of sowing (TOS including late winter, spring and summer) or depth by TOS by irrigation on the surface area, root dry weight, and specific root length in the 2019-2020 season (a, b and c, respectively) and 2020-2021 season (d, e, and f, respectively) at the flag leaf stage, for a sorghum crop (Genotype E) sown at 9 pl m^-2^ at Nangwee, Queensland, Australia. Values were the estimated means. Error bars represent standard errors of the estimations.

In the 2020-2021 season, summer sown sorghum had a significantly larger root volume (Fig. 2h, Table 5) compared to the spring and late winter sorghum, particularly in the first 0.3 m soil depth. It was also the case for the average root diameter (Fig. 2g) at soil depths above 1.3 m. However, there were no significant effects of TOS, irrigation, or their interactions with depth observed on the other root traits (Table 5).

**Table 5.** Results (p values) from the linear mixed model analysis on the root traits at the across-depth level for the main effects including depth (0 – 2.1 m with 8 depth intervals), time of sowing (TOS, late winter, spring, and summer), and irrigation (dryland and irrigation) and their interactions for a sorghum crop (Genotype E) sown at 9 pl m^-2^ in the 2019-2020 and 2020-2021 seasons at Nangwee, Queensland, Australia.

| Sources of variations | Root length<br>(cm) | Root<br>dry<br>weight<br>(g) | Specific<br>root<br>length<br>(cm g<sup>-1</sup>) | Surface<br>area<br>(cm<sup>2</sup>) | Root<br>volume<br>(cm<sup>3</sup>) | Root<br>length<br>density<br>(cm<br>cm<sup>-3</sup>) | Average<br>root<br>diameter<br>(cm) |
| --- | --- | --- | --- | --- | --- | --- | --- |
| **2019-2020** |  |  |  |  |  |  |  |
| Depth | 0.00*** | 0.000*** | 0.658 | 0.000*** | 0.000*** | 0.000*** | 0.000*** |
| TOS | 0.741 | 0.819 | 0.025* | 0.240 | 0.278 | 0.992 | 0.002** |
| Irrigation | 0.225 | 0.480 | 0.673 | 0.366 | 0.189 | 0.186 | 0.240 |
| Depth: TOS | 0.002** | 0.645 | 0.400 | 0.085 | 0.124 | 0.009** | 0.047* |
| Depth: Irrigation | 0.245 | 0.691 | 0.368 | 0.274 | 0.795 | 0.032* | 0.550 |
| TOS: Irrigation | 0.391 | 0.912 | 0.475 | 0.147 | 0.111 | 0.699 | 0.029* |
| Depth: TOS: Irrigation | 0.193 | 0.260 | 0.076 | 0.136 | 0.026* | 0.198 | 0.082 |
| **2020-2021** |  |  |  |  |  |  |  |
| Depth | 0.000*** | 0.000*** | 0.166 | 0.000*** | 0.000*** | 0.000*** | 0.000*** |
| TOS | 0.879 | 0.435 | 0.297 | 0.631 | 0.806 | 0.912 | 0.417 |
| Irrigation | 0.949 | 0.313 | 0.602 | 0.674 | 0.876 | 0.805 | 0.415 |
| Depth: TOS | 0.371 | 0.073 | 0.756 | 0.274 | 0.020* | 0.833 | 0.410 |
| Depth: Irrigation | 0.881 | 0.445 | 0.154 | 0.609 | 0.085 | 0.874 | 0.071 |
| TOS: Irrigation | 0.517 | 0.577 | 0.848 | 0.221 | 0.206 | 0.609 | 0.444 |
| Depth: TOS: Irrigation | 0.225 | 0.500 | 0.447 | 0.692 | 0.525 | 0.473 | 0.021* |

Where \*, \*\* and \*\*\* represents the significant level $\leq 0.05$ , $\leq 0.01$ and $\leq 0.001$ , respectively.

The root activity factor (R) calculated in eq. 2 (as in Zhao et al., 2022), was linearly related to the measured root length density (RLD) (Fig. 4), this is the larger the root length density the larger the root activity factor. Fig. 4 also shows that for similar values of root length density, the dryland plots had a larger values root activity than that the irrigated plots.

**Figure 4.**
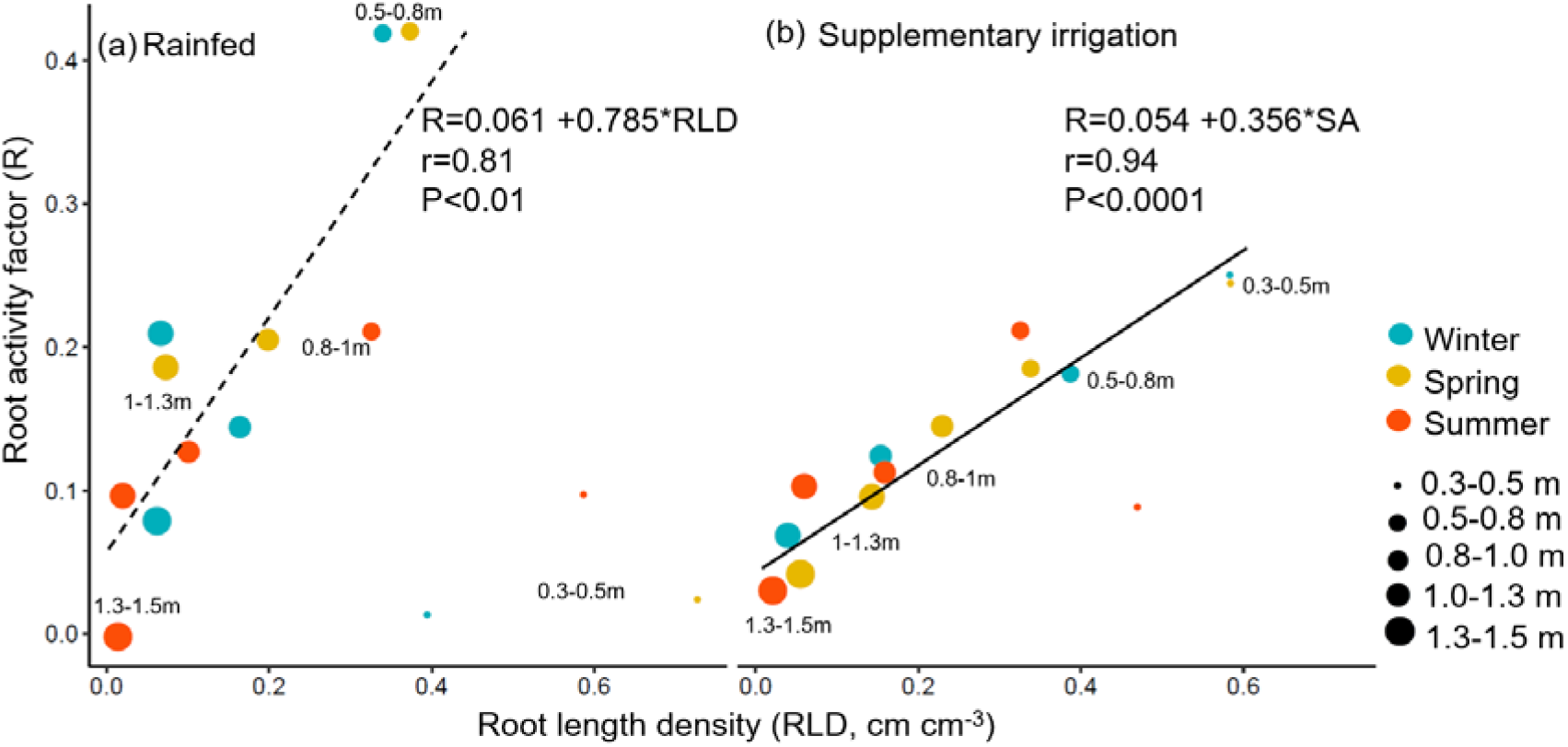
Relationship between the root activity factor (R) at flowering and root length density (RLD, cm^3^ cm^-3^) for the rainfed/dryland (a) and irrigated (b) plots. The data is for genotype E, sown at 9 pl m^-2^ in the 2020-2021 season. Blue, orange, and red dots indicate the late winter, spring and summer sown crops, respectively, and the size of the points indicate the soil layer. The linear relationships were not fitted to the data from the 0-0.3m and 0.3-0.5m depths, as those layers were close to wilting point, particularly in the rainfed treatment.

### 3.3. Soil water dynamics

In the first season (Fig. S2a), differences in PAW between TOS were small. During the second season (Fig. S2b) PAW values were much larger with larger differences between TOS. For example, soil moisture at the sowing of the late winter sown crop was 175 mm compared to that of spring (201 mm) and summer (253 mm) sown crops. At flowering, the irrigated late winter sown plots had larger values of PAW (272 mm) than the spring (211 mm) and summer (211 mm), while in the dryland plots PAW was similar across TOS (∼180 mm). At around 50 days after sowing, the smaller plant populations had larger values of PAW regardless of the TOS. In addition, more water was left in the soil profiles in the irrigated late winter and spring sown crops at the end of the season. The effect was more pronounced for the smaller plant populations.

The cumulative crop water use from sowing to flowering, from flowering to maturity, and from sowing to maturity (total water use) was also derived from the EMI surveys. In the 2019-2020 season, the difference in water use between TOS was small due to the dryness of the season (Fig. 5a-c). However, in the 2020-2021 season, sowing dates significantly changed sorghum water use dynamics around the crop critical stages. Fig. 5d shows that the late winter sown crop had the smallest pre-flowering water use at both irrigation levels with the spring having the next smallest values. In contrast, the summer sown crops had the largest pre-flowering water use (Fig. S2b). However, the opposite was true for the post-flowering water use (Fig. 5e) in which the late winter sown crop had the largest values, followed by the summer and spring sown crops. Summer sown crops had the largest values of total water use (Fig. 5f), followed by the spring and late winter sown crops, in that order, regardless of irrigation levels.

**Fig. 5.**
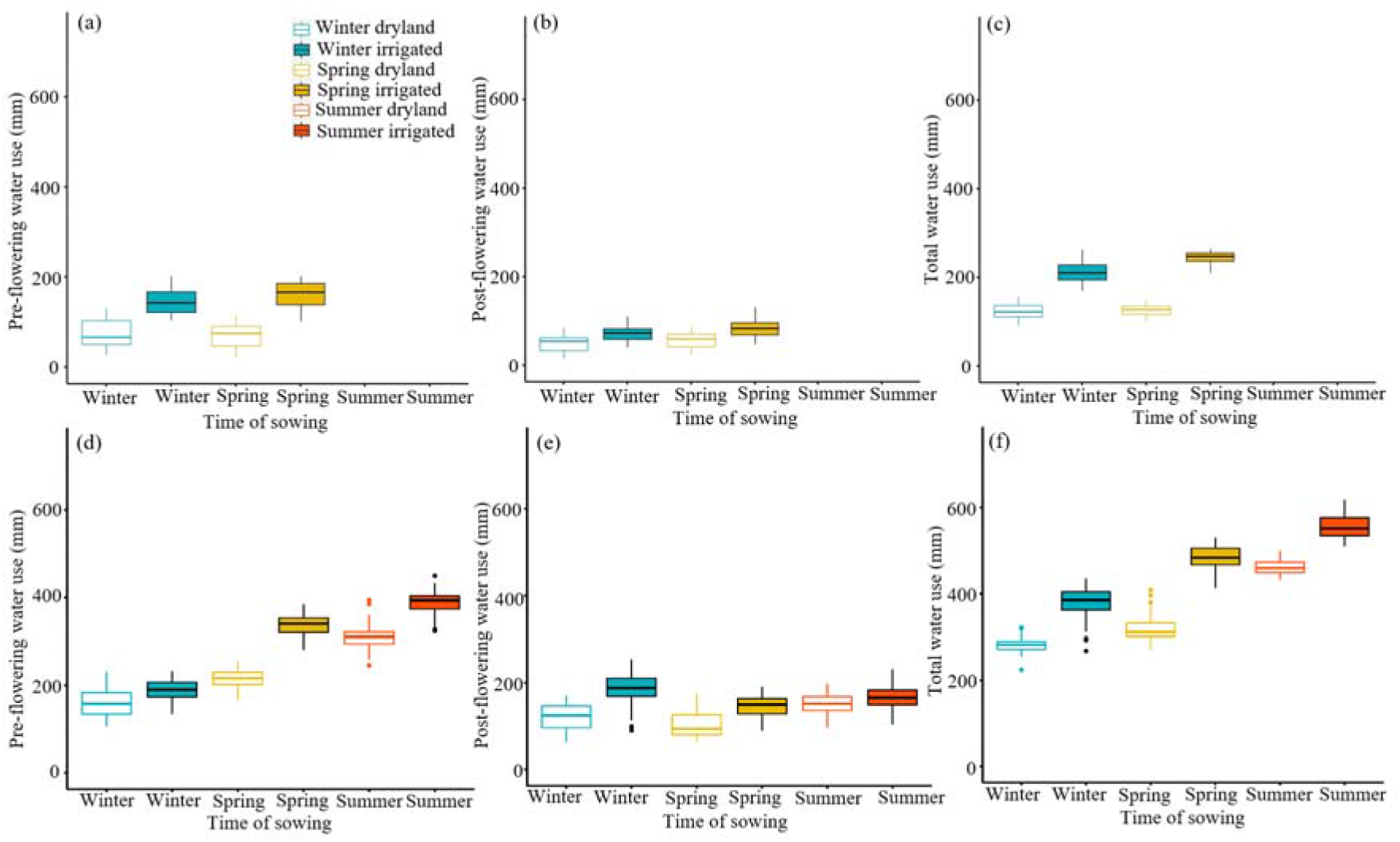
Cumulative crop water use (mm) derived from the electromagnetic induction surveys at (a) pre-flowering, (b) post-flowering, and (c) whole growing season in the 2019-2020 season and at (d) pre-flowering, (e) post-flowering, and (f) whole growing season in the 2020-2021 season at Nangwee, Queensland, Australia. Box and whisker plots in (a) – (f) show the median (central line), 25^th^ and 75^th^ percentiles (box), and 10^th^ and 90^th^ percentiles (whisker) of the crop water use.

### 3.4. Yields and yield components

The grain yield in the second season (4.05-7.03 t ha^-1^) was significantly larger than in the first season (0.95-1.62 t ha^-1^). In both seasons, spring sown sorghum had larger or similar yields than the late winter sown crop, and higher than the summer sown sorghum, especially in the irrigated treatments (Fig. 6a). This was associated with larger grain numbers for the late winter and spring sown crops, and similar or slightly lower grain weights (Fig. 6b and c).

**Fig. 6.**
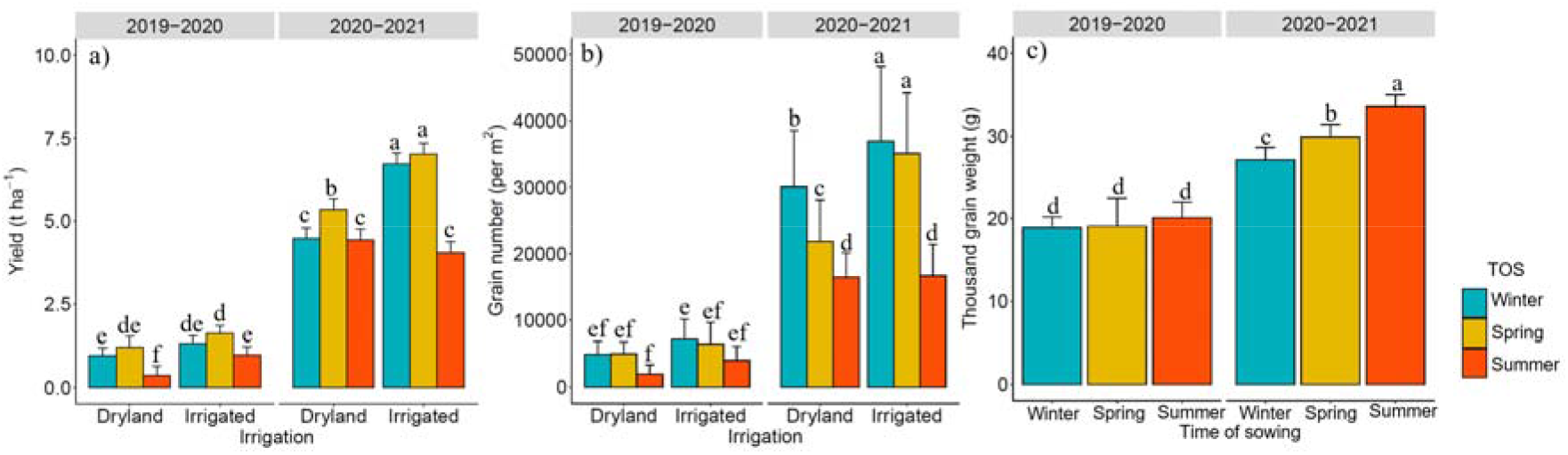
Effects of time of sowing (TOS, i.e., late winter, spring, and summer) or TOS by irrigation (i.e., dryland and irrigation) on the (a) grain yield, (b) grain number, and (c) grain weight across the 2019-2020 and 2020-2021 seasons at Nangwee, Queensland, Australia. Different lowercase letters indicate a significant difference at *p* ≤ 0.05. Error bars represent standard errors of the estimations.

As expected, strong positive relationships were observed between the grain yield and grain number (Fig. 7 and 8). Fig. 7b, c and Fig. 8b, c further indicate that the larger yield of early sown sorghum was contributed by the higher grain numbers, particularly in tillers. Irrespective of the origin (tillers or main stem) there was no relationship between grain yield and grain weights (Fig. 7d-f, Fig. 8d-f).

**Fig. 7.**
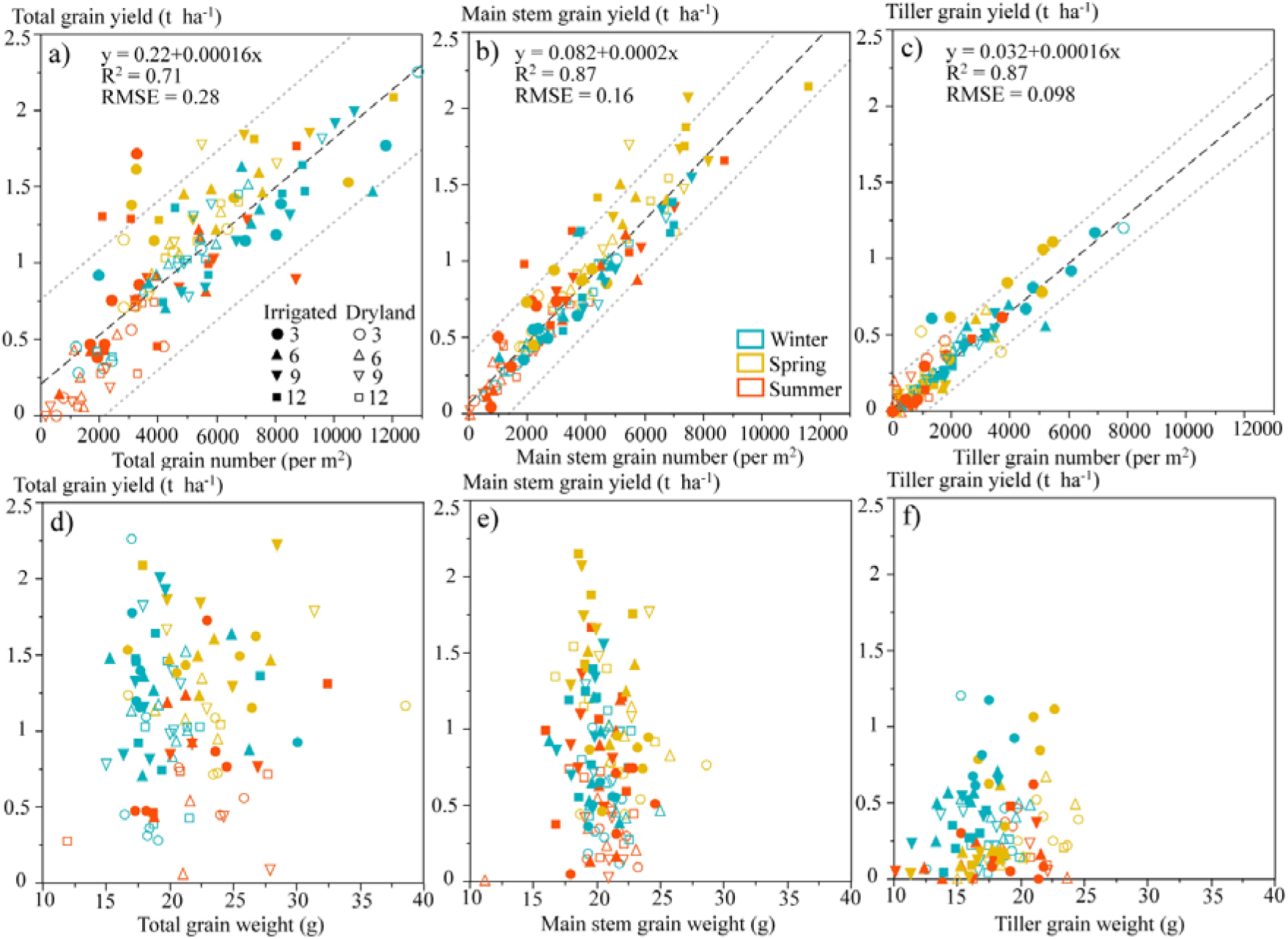
Relationship, for the 2019-2020 season, between sorghum grain yield and total grain number (a), main stem yield and grain number (b), tiller yield and grain number (c), and between yield and grain weight (d), main stem grain yield and grain weight (e), and tiller yield and grain weights (f). The fitted line and the associated 95% prediction interval are given as dashed lines and dotted lines, respectively. Values were estimated means for the three replicates from the linear mixed model as indicated in Table S3-S10.

**Fig. 8.**
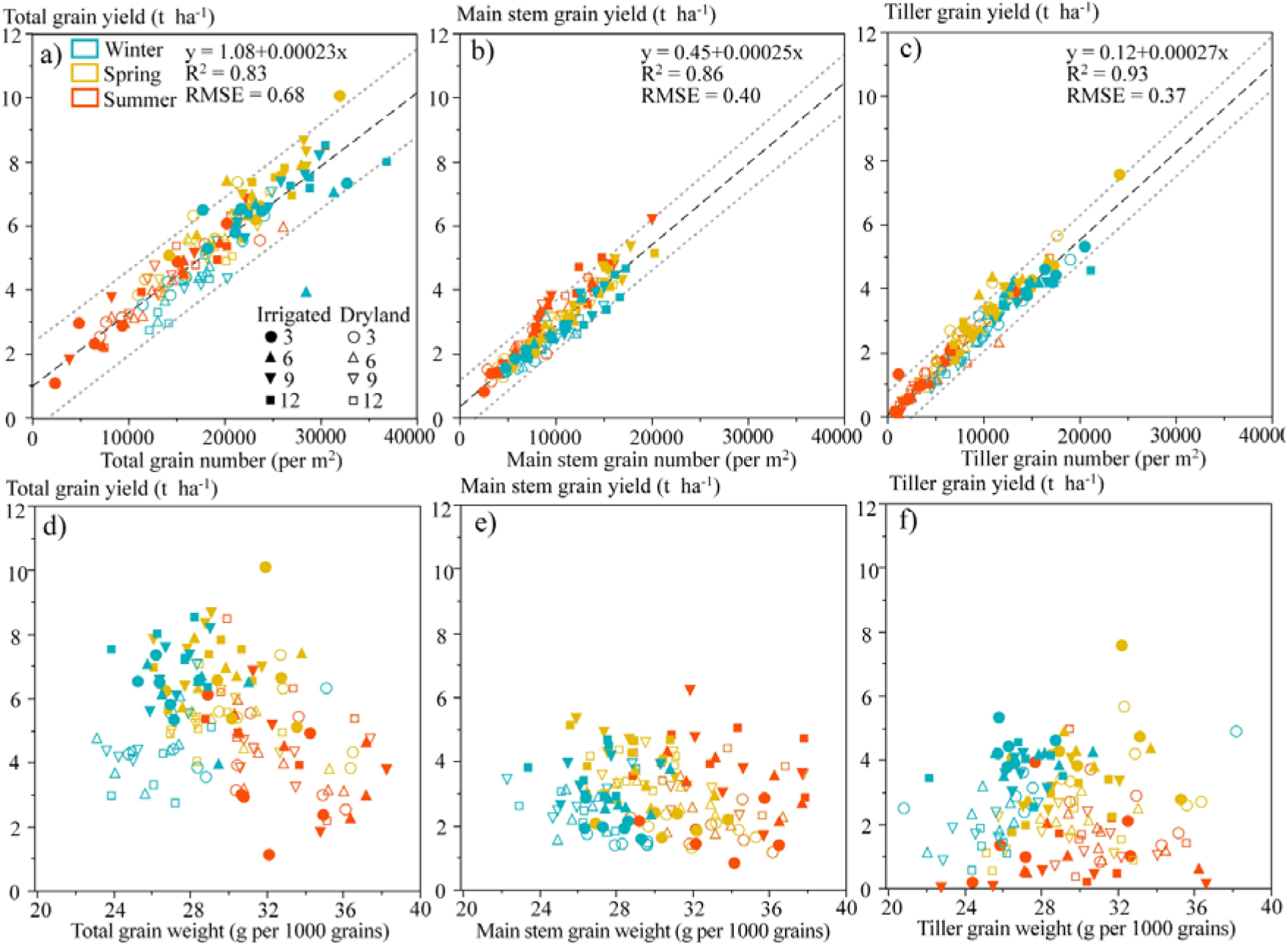
Relationship, for the 2020-2021 season, between sorghum grain yield and total grain number (a), main stem yield and grain number (b), tiller yield and grain number (c), and between yield and grain weight (d), main stem grain yield and grain weight (e), and tiller yield and grain weights (f). The fitted line and the associated 95% prediction interval are given as dashed lines and dotted lines, respectively. Values of each point were estimated means for the three replicates from the linear mixed model as indicated in Table S3-S10.

Irrigated treatments had significantly larger yields and grain numbers for both main stem and tillers (Fig. S3a, g). The yield and grain number for main stem increased with the increase in plant population densities, while, as expected, the opposite was true for tillers (Fig. S3b, h). Grain weight on tillers decreased with the increase in population densities (Fig. S3j), while the effect of population densities on main stem grain weight was inconsistent (Fig. S3e).

### 3.5. Relationships between root traits, water use, yield components, and environments

In the 2019-2020 season (Fig. 9a), PC1 explained 45.7% of variations in the dataset, which could largely be attributed to the environmental variables and root traits, while PC2 explained 21.2% of variations which can be attributed to the yield components, biomass, and water use. As expected, the points corresponding to late winter, spring, and summer sorghum were separated from each other. The magnitude and direction of the variables were different among TOS. In general, the early sown crops had a larger yield (i.e., grain yield and grain number) and harvest index values, which was associated with a higher value of post-flowering water use and water use efficiency (WUE).

**Fig. 9.**
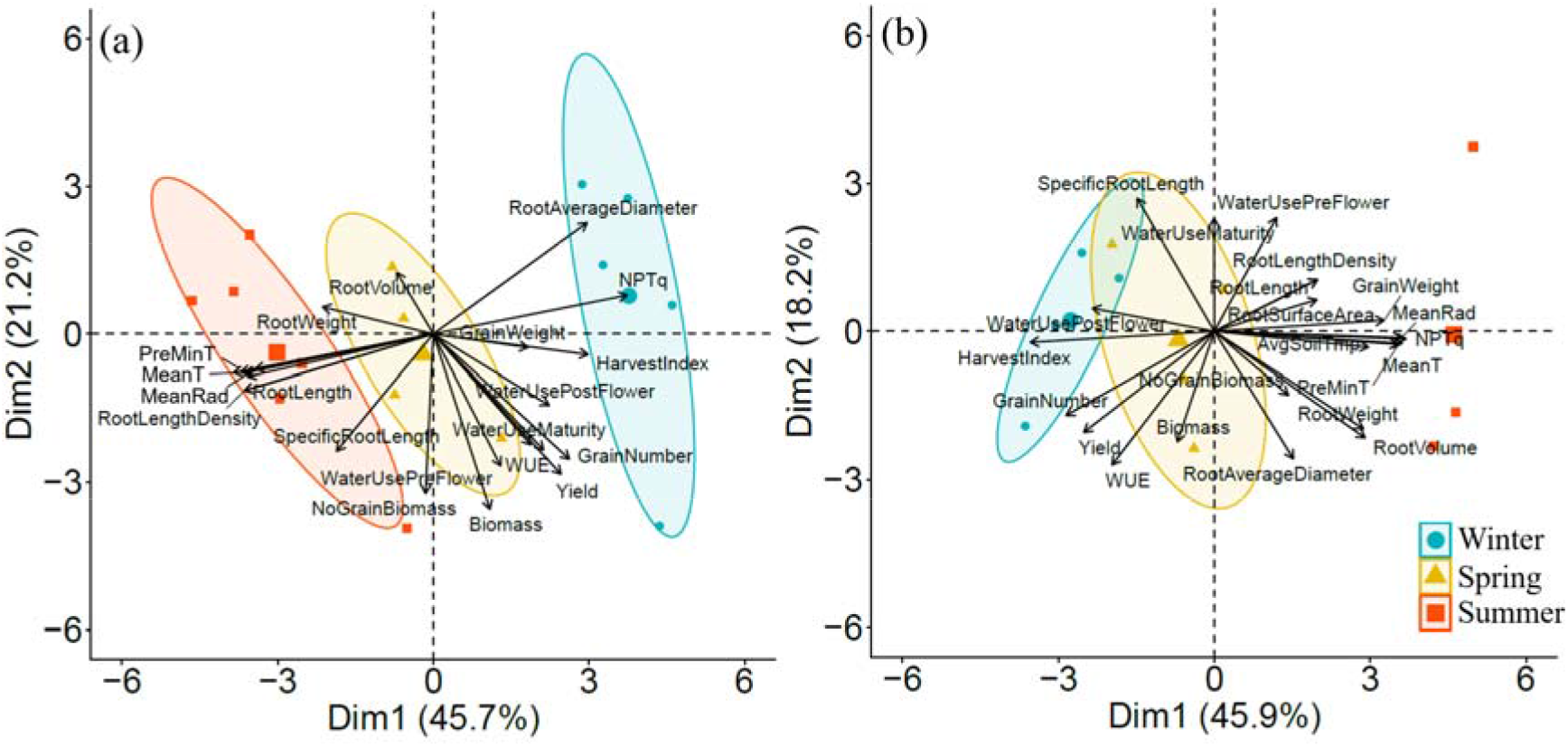
Principal component analysis of root traits, biomass, harvest index, yield components (i.e., yield, grain number, and grain weight), crop water use (i.e., pre-flowering water use – WaterUsePreFlower, post-flowering water use – WaterUsePostFlower, total water use – WaterUseMaturity) and environmental variables (i.e., mean temperature – MeanT, mean pre-flowering minimum temperature – PreFlwMinT, mean radiation – MeanRad, Normalised photo-thermal quotient – NPTq) across the (a) 2019-2020 and (b) 2020-2021 seasons at Nangwee, Queensland, Australia. Each time of sowing was identified by a different symbol and a 68% confidence limit ellipse.

Fig. 10a shows a positive linear relationship between WUE and the total grain number across both seasons for the irrigated and dryland treatment, respectively. Particularly, larger values of WUE in the late winter sown crops were related to a larger grain number from tillers (Fig. 10b). In addition, Fig. 10c shows WUE increased with the ratio of post-flowering water use to total water use, with the effects at dryland being more significant than the irrigated.

**Fig. 10.**
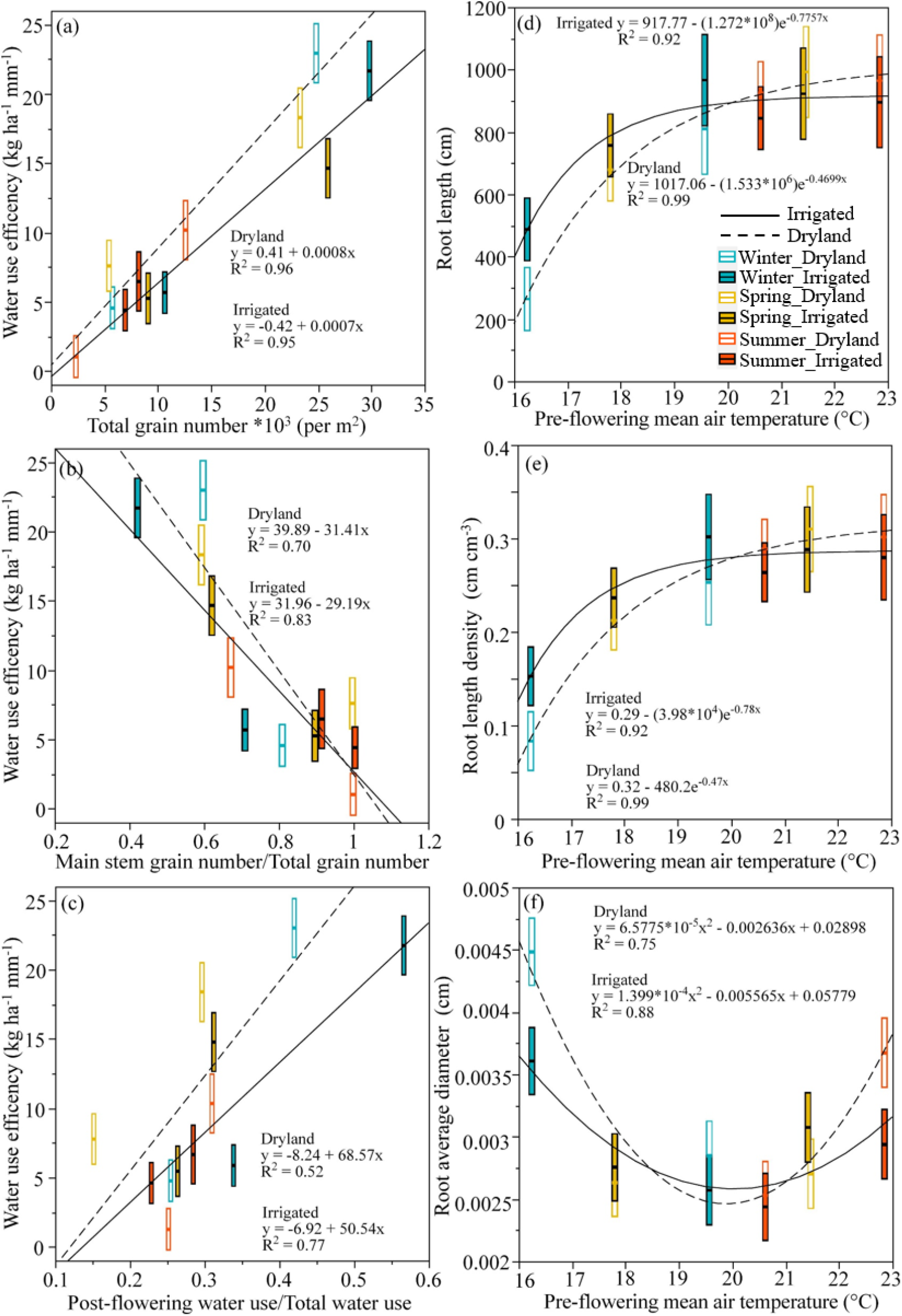
Relationships between (a) water use efficiency (kg ha^-1^ mm^-1^) and total grain number (per m^2^), (b) the ratio of main stem grain number to total grain number and (c) ratio of post-flowering water use to total water use and between plot-level (d) root length (cm), (e) root average diameter (cm) and (f) root length density (cm cm^-3^) and pre-flowering mean air temperature (°C), respectively, across the two study seasons for the three times of sowing (i.e., late winter, spring and summer) and two water levels (i.e., dryland and irrigated). The horizontal line in each box is the estimated mean with upper and lower bounds for standard errors of the estimation.

Fig. 9a also shows an association between the root length, root weight, root surface area, root length density, and specific root length of summer sown sorghum and ambient temperature and solar radiation co-variates. This is shown in Fig. 10d which shows the relationships between pre-flowering mean air temperature and root length across seasons and shows that the increase in root length with increasing pre-flowering mean air temperature is exponential, according to the equations y= 1017.06 - 1.533 * 10^6^e^-0.4699x^ and y = 917.77 - 1.272 * 10^8^e^-0.7757x^ for the dryland and irrigated, respectively. The maximum values for the root length were observed between 19 to 20 °C for the irrigated, and >23 °C for the dryland treatments. Similarly, root length density also followed exponential trends with pre-flowering mean air temperature for both irrigation levels (Fig. 10e).

In addition, in the 2019-2020 (cooler) season, the late winter sown sorghum had the largest average root diameter (thicker roots), though this was not observed in 2020-2021 (warmer) season (Fig. 9b). Root average diameter fitted quadratic relationships with pre-flowering mean air temperature (Fig. 10f), with the smallest average root diameter values observed around 20 °C, irrespective of irrigation treatment.

Results from the 2020-2021 season (Fig. 9b), show that PC1 explained 45.9% of variations in the dataset, while PC2 explained 18.2% of variations, like the first season, with a high loading for the root traits and yield components. Missing values for some of the yield components in summer sown crops didn’t allow us to calculate the ellipse in normal probability. Like the 2019-2020 season, larger values of harvest index, yield, and grain number were observed for the late winter and spring sown crops, which were associated with larger values of WUE. The summer sown sorghum had a larger rooting system which was positively associated with larger temperature and solar radiation co-variates and pre-flowering water use.

## 4. Discussion

The tested sowing dates across seasons significantly changed the temperature, radiation, and hydric environments of the crops. The early sown crops decreased the risk of extreme heat around flowering and increased the likelihood of higher grain numbers i.e., higher seed set values. Sowing sorghum, a summer crop, in late winter into cold soils and chilling ambient temperatures altered root growth, biomass production, water use dynamics, and yield components. Yield differences between late winter and spring sowing were small due to important compensations between main stem and tillers i.e., a larger grain number contribution from tillers in the late winter sown crop. Sowing sorghum in late winter had negative effects on root growth leading to a smaller crop with a smaller rooting system and thicker roots. More water was left in the soil profile at the end of the season after a late winter sown crop, particularly under irrigation and low plant populations.

### 4.1. Crop phenology

As expected, lower ambient temperatures for the late winter and spring sown crops delayed crop emergence and flowering, though still the late winter sown crop flowered earlier than the spring and summer alternatives reducing the likelihood of heat stresses around flowering (Kapanigowda et al. 2013; Barbosa et al., 2020). This was also reflected in lower values for the estimated seed set (Singh, et al., 2015).

### 4.2. Crop growth and root growth and function

Chilling stresses early in the growing season have been shown to affect metabolism and photosynthesis, including thylakoid electron transport, carbon assimilation, and stomatal control (Abbas, 2012; Bekele et al., 2014; Casto et al., 2021). Cold soil temperatures reduce root growth by reducing the availability of sugars and root development by initiating second and third-order laterals and inhibition of root elongation (Kaspar and Bland, 1992; Nagel et al., 2009). Our results also showed that late winter sowing produced thicker roots significantly reducing total root length, root length density and root volume, compared to the spring and summer sowing. As shown in maize (Farooq et al., 2009; Zhou et al., 2021) and wheat (Miyasaka and Grunes, 1990; Hassan et al., 2021), cold temperatures significantly increased the root diameter of sorghum. This could be attributed to the inhibition of root branching (Clowes and Wadekar, 1988). Possible explanations for the limited root branching and thicker roots are that low temperatures on the root meristems might affect the production of growth substances, and or reduce the uptake of diffusion affected nutrients such as potassium and phosphorus (Koevoets et al., 2016; Zhou et al., 2022). The root activity factor was not calculated during the first season of trials due the lack of available EMI surveys around flowering (Fig. S1a). For the second season of trials, the calculated root activity factor was closely related to root length density across most of the soil profile. The decline in the root activity factor with soil in-depth was previously related to a lack of time for the rooting system to explore deeper soil layers by flowering (Robertson et al., 1993). In general, the larger the root length density the larger the values of the root activity factor (Figs. 4). However, different relationships were evident for the irrigated and dryland treatments. For the same value of root length density, dryland plots had larger values of the root activity factor than the irrigated plots. In the top layer of the dryland plots root activity was limited by water supply irrespective of the root length density. In the irrigated plots, the slope for the relationship between RLD and R was larger for the summer sown sorghum crops (like the dryland plots) compared to the late winter and Spring sown crops. The differences in slope between the dryland and irrigated and summer sown and early sown might be related to a stress adaptation (e.g., an increase in root hair and length in water-limited environment) (Calleja-Cabrera et al., 2020), and or differences in atmospheric demand (Robertson et al., 1993), respectively.

We showed (Fig. 10) that root length and root length density were affected by pre-flowering mean temperatures. Both traits followed a typical temperature response curve in which the root length increased with the increasing temperature to an optimal temperature of 25-30 °C (Kaspar and Bland, 1992). Here we propose that root growth and branching in cold soils might be an important trait to include in the selection for more cold-tolerant sorghum genotypes.

### 4.3. Soil water dynamics and crop yield determination

Sowing dates affected crop water use dynamics differently across seasons. Small differences were observed between treatments in the drier and cooler first season of trials. In the second, warmer and wetter season, the late winter sown sorghum had the smallest pre-flowering water use (Fig. 5), which was associated (Fig. 9) with the smaller size of the rooting system and reduced atmospheric demand (Table 2). The reduced pre-flowering water use and the relatively higher grain numbers and grain yields (Fig. 6) produced higher water use efficiency values for the late winter and spring sown sorghum compared to the summer sown crop (Carcedo et al., 2021). The difference in post-flowering water use due to the time of sowing was responsible for an increased yield, grain number, biomass, and harvest index of early sown sorghum. Compensations between grain number and grain yield are common in cereals (Gambín and Borrás, 2007). In the case of sorghum, phenotypic plasticity (Sadras and Slafer, 2012) was also observed with the increased production of tillers and the contribution of tillers to total grain number in the late winter sown crops (Clarke, et al., 2019). These results align with the observations by Rodriguez et al., (2023 – this special issue) that the avoidance of heat stress and the transfer of water use from vegetative to reproductive stages in late winter sown sorghum are key factors in determining the final yield of early-sown sorghum. In addition, larger residual moisture was observed particularly for the irrigated early sowing, and lower plant populations. Larger values of residual moisture can increase the chances of double cropping a winter crop after a short summer fallow. The larger pre-flowering water use of the summer sorghum reduced post-flowering water availability. Thus, sowing date affected the dynamics of water stress and the resilience of dryland cropping systems (Hammer et al. 2014; Rodriguez et al. 2018; Rodriguez et al., this special issue). An understanding of the impact of time of sowing on soil and crop water status can help define management strategies to increase crop productivity (Clarke et al., 2019).

### 4.4. Terminal heat avoidance and early-stage cold tolerance as a target

Ongoing climate change can be expected to increase the intensity and frequency of heat waves and drought events (IPCC, 2021). In sorghum, a 10-15-day window centred around flowering is the most sensitive phase of sorghum to heat waves (Prasad et al., 2008; Singh et al., 2017). High-temperature stresses during this window will increase pollen sterility and reduce seed set, causing grain yield losses (Singh et al., 2016, 2017). Adaptation strategies are required to sustain sorghum production and maintain profits of the yield under climate change (Kothari et al., 2020). In the long term, breeding can be expected to contribute to improving genetic tolerance to heat stresses (Singh et al., 2014), while agronomy could immediately increase adaptation by avoiding the overlap between times of the year with the high frequency of heat stress events and flowering windows (Kothari et al., 2020). Plant breeders have been already targeting plant traits to develop elite cultivars more adaptable to high temperatures (Nguyen et al., 2013; Tack et al., 2017; Janni et al., 2020). However, the complex multi-genic control of associated traits and complicated G × E scenarios can limit progress (Jha et al., 2014). Within this context, early sowing can play an important role in improving crop adaptability to future climates before well-adapted cultivars are released (Munaro et al., 2020). Early sowing allows the crop to flower before heat waves and mature before the terminal drought, subsequently enhancing productivity (Chiluwal et al., 2018; Vennapusa et al., 2021). Moreover, it can also improve the use efficiency of soil residual water and reduce evaporative soil water losses during later stages (Hund et al., 2009). This is further evidenced by the present study in which a significantly higher yield, seed set, and WUE were achieved by the early-sown sorghum compared to the summer-sown crop.

However, sorghum is highly sensitive to cold temperatures (Rooney, 2004) requiring soil bed temperatures higher than 18 °C for germination and seedling establishment (Shroyer et al., 1998; Ostmeyer et al., 2020). In this study, cold soil temperatures at the early growing stages significantly limited root growth and development of late winter sown crops. This led to a smaller root and shoot system, which negatively affected the yield in comparison to the spring sown crops. Sorghum crops exposed to consecutive cold stress and excessive light conditions have a significantly lower rate of photosynthetic CO_2_ uptake (Ortiz et al., 2017), affecting the dynamics of crop growth and water use.

Previous studies have also identified negative effects of early-stage cold stress on germination (Upadhyaya et al., 2016), seedling emergence and survival (Parra-Londono et al., 2018), and seedling vigor (Moghimi et al., 2019). Chiluwal et al. (2018) reported that early sowing resulted in significantly smaller leaf area, shoot biomass, and root system than regular sowing. Similarly, Moghimi et al. (2019) observed a delay in germination and emergence and ∼61% and 73% lower root and total seedling biomass, respectively, for the sorghum under cold stress compared to the normal crop. Recently, Ostmeyer et al. (2020) reported significantly smaller seedling shoot-root ratios for the early sowing sorghum than regular sowings.

Based on these results, we propose that in the race to increase sorghum adaptation to heat stress breeding should seriously consider cold tolerance during crop germination, emergence, and vegetative stages as a target, so that sowing dates could be significantly advanced without affecting the crop’s establishment and early growth. Recent studies have identified promising candidate genes putatively conferring germination (Upadhyaya et al., 2016), seeding emergence and survival (Parra-Londono et al., 2018), and seedling vigour with active photosynthesis and chlorophyll retention (Moghimi et al., 2019; Vennapusa et al., 2021) cold tolerance. Related studies suggest that under cold stress the development of the crop root system controls the initial success or failure of the seedling (Enns et al., 2006; Farooq et al., 2009). For instance, an increased lateral root development could contribute to greater cold tolerance (Hund et al., 2009; Chopra et al., 2017). Thus, the variability of the length of lateral roots could be exploited to improve sorghum growth at the early growing stages. In addition, antioxidant capacity increases during cold acclimation in several plants as an adaptive mechanism to low temperatures (Foyer and Noctor 2005).

## 5. Conclusion

Root growth appears to be negatively affected in early sown crops (smaller root systems). Sowing sorghum, a summer crop, in late winter reduced water use during vegetative stages and transfer water use between flowering and maturity. This produced higher grain weights, and grain numbers in tillers. Early sown sorghum is more likely to avoid heat stresses around flowering increasing the likelihood of higher grain numbers and grain yields in the hottest seasons. In the race to increase crop adaptation early sowing of summer crops like sorghum is possible, though breeders should seriously consider cold tolerance during crop germination, emergence, and vegetative stages as a target so that the risk of poor crop establishment and growth early in the season due to cold conditions is minimised.

## Acknowledgments

This work is funded by the Australian Grains Research and Development Corporation project (GRDC UOQ1906-010RTX).

## CRediT authorship contribution statement

**Dongxue Zhao:** Formal Analysis, Writing - Original Draft, Writing - Review & Editing; **Daniel Rodriguez**: Project leadership, Conceptualization, Methodology, Formal Analysis, Writing & Editing; **Peter deVoil**: Data management, Review & Editing; **Bethany Rognoni and Kerry Bell:** statistical data analysis & Editing; the rest of the authors contributed by running field experiments, collecting data and review.

## Data availability

The data is available upon request to the corresponding author and approval from the funding body (GRDC).

