## Supplemental materials for "Adapting to heat stress by sowing summer grain crops early in late winter: Sorghum root growth, water use, and yield"

Supplementary information associated with:

**Adapting to heat stresses by sowing summer grain crops in winter: Sorghum root growth and function in cold soils**

Dongxue Zhao et al., 2023

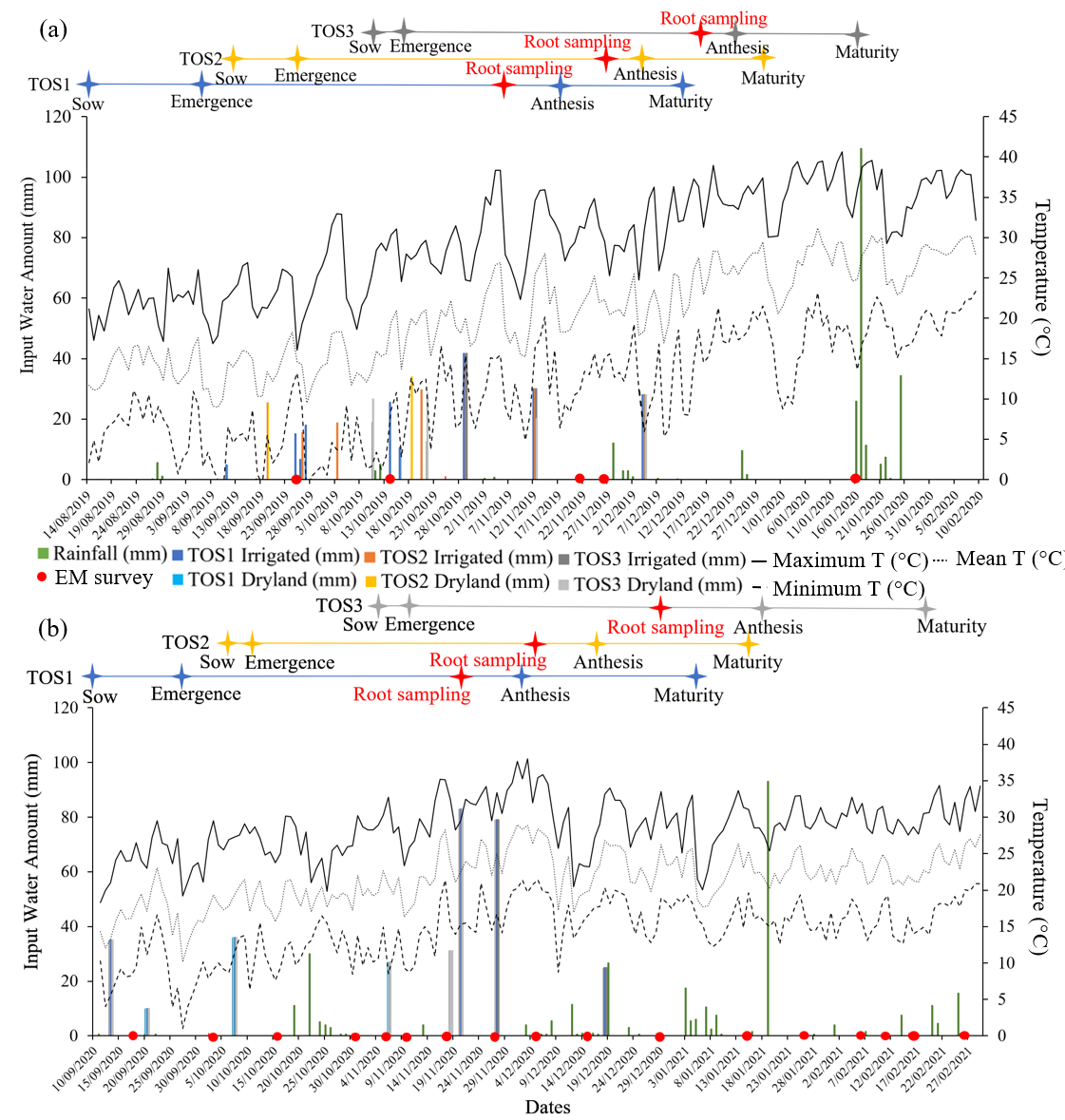

**Figure S1.** Crop phenological stages for the three times of sowing (TOS, i.e., late winter-TOS1, spring-TOS2 and summer-TOS3) and dates of the electromagnetic induction (EMI) surveys; rainfall and irrigation amounts; and maximum, minimum, and mean temperature (T) are indicated in the (a) 2019-2020 and (b) 2020-2021 Southern Hemisphere summer growing seasons. *The phenology stage for each TOS is the mean value for all the treatments including six genotypes, two irrigation levels, and four plant populations.

| **Table S1.** The sorghum phenology for the three times of sowing (TOS) includes late winter, spring and summer, two levels of irrigation, and six commercial genotypes at 2019-2020 and 2020-2021 growing seasons at Nangwee, Queensland, Australia. Values indicate days or cumulative degree days from sowing to emergence, emergence to flowering and emergence to maturity, respectively. Ranges indicate variability between treatments other than TOS. | | | | | | | | | | | | | | | |
| --- | --- | --- | --- | --- | --- | --- | --- | --- | --- | --- | --- | --- | --- | --- | --- |
|  |  |  | 2019-2020 | | | | | |  | 2020-2021 | | | | | |
|  |  |  | Emergence | | Emergence-Flowering | | Emergence-Maturity | |  | Emergence | | Emergence-Flowering | | Emergence-Maturity | |
| TOS | Irrigation | Genotype | Days | CGD  (°C d) | Days | CGD  (°C d) | Days | CGD  (°C d) |  | Days | CGD  (°C d) | Days | CGD  (°C d) | Days | CGD  (°C d) |
| Winter | Irrigated | A | 18 | 73 | 74 | 713 | 102 | 1165 | 173 | 8 | 52 | 76 | 787 | 109 | 1184 |
|  |  | B | 25 | 114 | 67 | 672 | 103 | 1261 | 176 | 9 | 59 | 71 | 708 | 108 | 1177 |
|  |  | C | 18 | 73 | 79 | 786 | 98 | 1086 | 176 | 8 | 52 | 72 | 715 | 109 | 1184 |
|  |  | D | 24 | 112 | 75 | 784 | 100 | 1199 | 175 | 9 | 59 | 75 | 780 | 108 | 1177 |
|  |  | E | 20 | 82 | 77 | 777 | 98 | 1115 | 173 | 9 | 59 | 71 | 708 | 108 | 1177 |
|  |  | F | 21 | 90 | 76 | 769 | 101 | 1182 | 174 | 9 | 59 | 75 | 780 | 108 | 1177 |
|  | Dryland | A | 19 | 77 | 78 | 782 | 105 | 1234 | 176 | 10 | 70 | 74 | 769 | 107 | 1166 |
|  |  | B | 28 | 122 | 69 | 737 | 98 | 1220 | 174 | 10 | 70 | 70 | 697 | 107 | 1166 |
|  |  | C | 22 | 98 | 77 | 798 | 104 | 1244 | 179 | 11 | 83 | 67 | 651 | 106 | 1153 |
|  |  | D | 20 | 82 | 77 | 777 | 108 | 1293 | 172 | 10 | 70 | 74 | 769 | 107 | 1166 |
|  |  | E | 23 | 106 | 74 | 753 | 101 | 1205 | 175 | 10 | 70 | 70 | 697 | 107 | 1166 |
|  |  | F | 27 | 120 | 72 | 776 | 103 | 1294 | 174 | 11 | 83 | 69 | 684 | 106 | 1153 |
| Spring | Irrigated | A | 14 | 119 | 63 | 764 | 93 | 1267 | 206 | 5 | 56 | 66 | 770 | 92 | 1066 |
|  |  | B | 17 | 152 | 60 | 731 | 94 | 1294 | 203 | 6 | 67 | 68 | 805 | 96 | 1119 |
|  |  | C | 15 | 130 | 64 | 789 | 96 | 1316 | 195 | 5 | 56 | 68 | 800 | 96 | 1114 |
|  |  | D | 15 | 130 | 68 | 847 | 96 | 1316 | 192 | 5 | 56 | 70 | 832 | 96 | 1114 |
|  |  | E | 15 | 130 | 62 | 753 | 92 | 1256 | 198 | 5 | 56 | 67 | 783 | 95 | 1100 |
|  |  | F | 15 | 130 | 68 | 847 | 96 | 1316 | 201 | 5 | 56 | 68 | 800 | 94 | 1087 |
|  | Dryland | A | 15 | 130 | 62 | 753 | 92 | 1256 | 204 | 5 | 56 | 66 | 770 | 92 | 1066 |
|  |  | B | 16 | 140 | 61 | 743 | 95 | 1306 | 202 | 5 | 56 | 68 | 800 | 96 | 1114 |
|  |  | C | 14 | 119 | 69 | 858 | 97 | 1327 | 206 | 5 | 56 | 67 | 783 | 96 | 1114 |
|  |  | D | 15 | 130 | 64 | 789 | 96 | 1316 | 199 | 5 | 56 | 69 | 816 | 95 | 1100 |
|  |  | E | 14 | 119 | 63 | 764 | 93 | 1267 | 200 | 5 | 56 | 68 | 800 | 96 | 1114 |
|  |  | F | 15 | 130 | 64 | 789 | 92 | 1256 | 206 | 5 | 56 | 67 | 783 | 95 | 1100 |
| Summer | Irrigated | A | 6 | 77 | 63 | 893 | 117 | 1799 | 239 | 5 | 46 | 55 | 714 | 106 | 1351 |
|  |  | B | 6 | 77 | 69 | 1004 | 117 | 1799 | 247 | 5 | 46 | 55 | 714 | 106 | 1351 |
|  |  | C | 6 | 77 | 69 | 1004 | 117 | 1799 | 244 | 5 | 46 | 55 | 714 | 106 | 1351 |
|  |  | D | 6 | 77 | 69 | 1004 | 117 | 1799 | 253 | 5 | 46 | 55 | 714 | 106 | 1351 |
|  |  | E | 6 | 77 | 63 | 893 | 117 | 1799 | 251 | 5 | 46 | 55 | 714 | 106 | 1351 |
|  |  | F | 6 | 77 | 69 | 1004 | 117 | 1799 | 244 | 5 | 46 | 55 | 714 | 106 | 1351 |
|  | Dryland | A | 6 | 77 | 63 | 893 | 117 | 1799 | 257 | 5 | 46 | 55 | 714 | 106 | 1351 |
|  |  | B | 6 | 77 | 63 | 893 | 117 | 1799 | 257 | 5 | 46 | 55 | 714 | 106 | 1351 |
|  |  | C | 6 | 77 | 69 | 1004 | 117 | 1799 | 262 | 5 | 46 | 55 | 714 | 106 | 1351 |
|  |  | D | 6 | 77 | 69 | 1004 | 117 | 1799 | 265 | 5 | 46 | 55 | 714 | 106 | 1351 |
|  |  | E | 6 | 77 | 63 | 893 | 117 | 1799 | 259 | 5 | 46 | 55 | 714 | 106 | 1351 |
|  |  | F | 6 | 77 | 69 | 1004 | 117 | 1799 | 257 | 5 | 46 | 55 | 714 | 106 | 1351 |
| CGD is cumulative growing degree days from sowing calculated using a base temperature of 10 °C and daily minimum and maximum temperatures from an automatic weather station installed in the field. | | | | | | | | | | | | | | | |

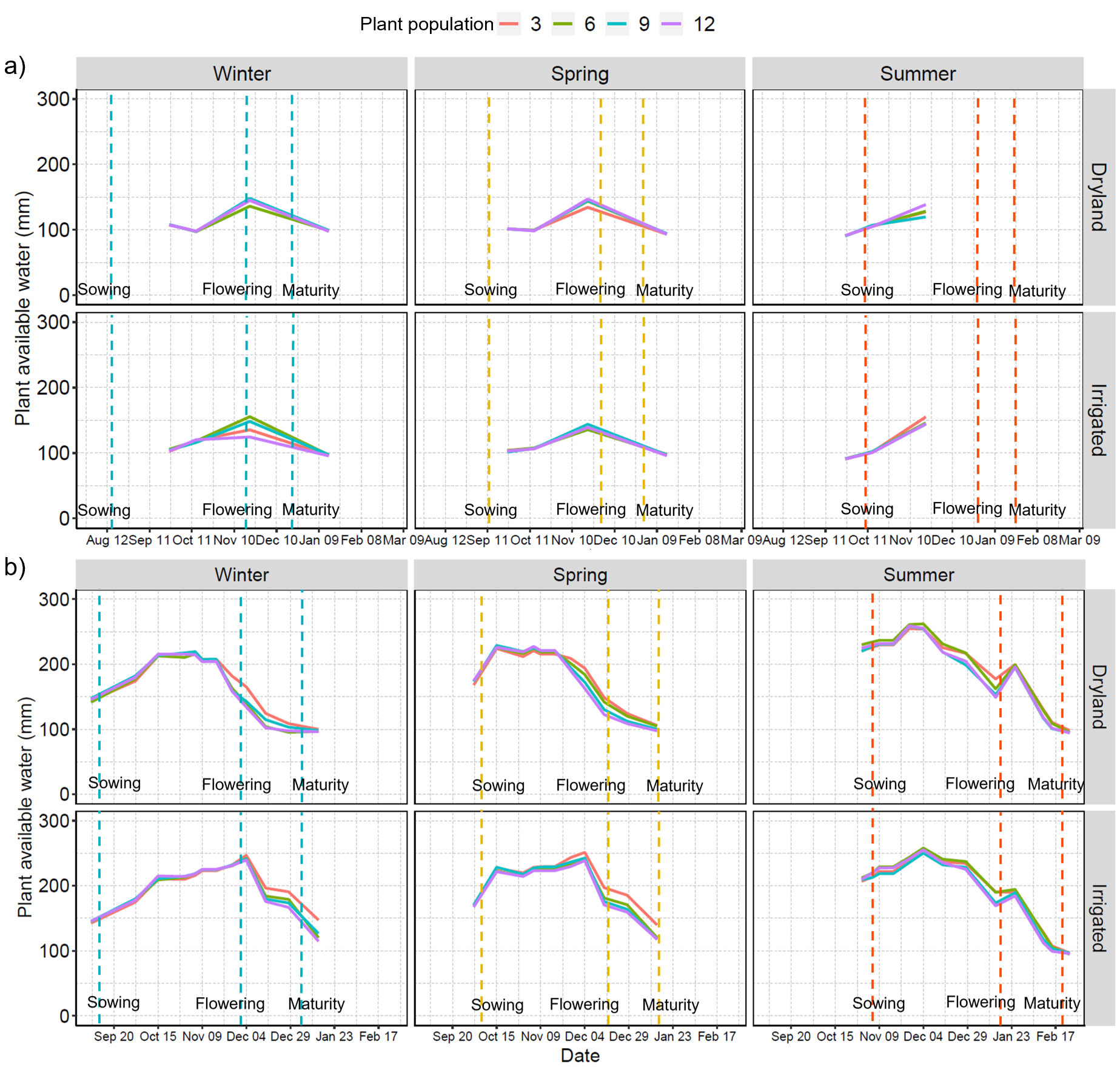

Figure S2. Plant available water content (mm) across the growing season derived from the electromagnetic induction (EMI) surveys in the (a) 2019-2020 and (b) 2020-2021 seasons at Nangwee, Queensland, Australia. Vertical dashed lines indicate the sowing, flowering, and maturity time.

| **Table S2.** Fixed effects testing results from the linear mixed model analysis of total grain yield for the late winter, spring, and summer sown sorghum in the 2019-2020 and 2020-2021 seasons using season, time of sowing (TOS), irrigation (Irri), plant population density (Pop) and genotype as fixed factors and replicates and main plot as random effects. | | | | |
| --- | --- | --- | --- | --- |
| Model parameter | Df | denDF | F-Value | *p*-Value |
| (Intercept) | 1 | 3.9 | 413.4 | 0.000^***^ |
| Season | 1 | 3.9 | 226.2 | 0.000^***^ |
| TOS | 2 | 14.8 | 29.5 | 0.000^***^ |
| Irri | 1 | 15.6 | 40.2 | 0.000^***^ |
| Pop | 3 | 230.7 | 5.4 | 0.001^**^ |
| Genotype | 5 | 234.4 | 13.5 | 0.000^***^ |
| Season: TOS | 2 | 15.6 | 8.3 | 0.004^**^ |
| Season: Irri | 1 | 16.0 | 6.5 | 0.021^*^ |
| TOS: Irri | 2 | 16.1 | 2.2 | 0.142 |
| Season: Pop | 3 | 304.5 | 1.9 | 0.136 |
| TOS: Pop | 6 | 246.3 | 1.7 | 0.128 |
| Irri: Pop | 3 | 253.9 | 1.1 | 0.338 |
| Season: Genotype | 5 | 316.6 | 4 | 0.002^**^ |
| TOS: Genotype | 10 | 248.3 | 2.4 | 0.011^*^ |
| Irri: Genotype | 5 | 253.5 | 0.8 | 0.588 |
| Pop: Genotype | 15 | 255.0 | 1.4 | 0.146 |
| Season: TOS: Irri | 2 | 16.5 | 12.3 | 0.000^***^ |
| Season: TOS: Pop | 6 | 314.6 | 0.9 | 0.498 |
| Season: Irri: Pop | 3 | 321.2 | 0.9 | 0.447 |
| TOS: Irri: Pop | 6 | 277.9 | 1.2 | 0.313 |
| Season: TOS: Genotype | 10 | 331.5 | 4.2 | 0.000^***^ |
| Season: Irri: Genotype | 5 | 334.1 | 1.4 | 0.233 |
| TOS: Irri: Genotype | 10 | 278.3 | 0.7 | 0.751 |
| Season: Pop: Genotype | 15 | 342.9 | 1.3 | 0.216 |
| TOS: Pop: Genotype | 30 | 266.8 | 0.6 | 0.931 |
| Irri: Pop: Genotype | 15 | 275.3 | 0.4 | 0.970 |
| Season: TOS: Irri: Pop | 6 | 332.1 | 2.5 | 0.020^*^ |
| Season: TOS: Irri: Genotype | 10 | 332.5 | 1.1 | 0.344 |
| Season: TOS: Pop: Genotype | 30 | 338.0 | 0.8 | 0.792 |
| Season: Irri: Pop: Genotype | 15 | 344.5 | 1.2 | 0.296 |
| TOS: Irri: Pop: Genotype | 30 | 306.9 | 0.4 | 0.997 |
| Season: TOS: Irri: Pop: Genotype | 27 | 340.1 | 2.1 | 0.002^**^ |
| Where ^*^, ^**^ and ^***^ represents significance at the ≤ 0.05, ≤ 0.01 and ≤ 0.001 levels, respectively. | | | | |

| **Table S3.** Fixed effects testing results from the linear mixed model fitted with residual maximum likelihood (REML) for predicting grain number for the late winter, spring, and summer sown sorghum in the 2019-2020 and 2020-2021 seasons using season, time of sowing (TOS), irrigation (Irri), plant population density (Pop) and genotype as fixed factors and replicates and main plots as random effects. | | | | |
| --- | --- | --- | --- | --- |
| Model parameter | Df | denDF | F-Value | *p*-Value |
| (Intercept) | 1 | 9.3 | 1505.0 | 0.000^***^ |
| Season | 1 | 3.8 | 302.8 | 0.000^***^ |
| TOS | 2 | 15.9 | 26.9 | 0.000^***^ |
| Irri | 1 | 14.7 | 31.2 | 0.000^***^ |
| Pop | 3 | 182.9 | 4.4 | 0.005^**^ |
| Genotype | 5 | 181.7 | 12.2 | 0.000^***^ |
| Season: TOS | 2 | 18.8 | 4.1 | 0.034^*^ |
| Season: Irri | 1 | 18.4 | 2.4 | 0.136 |
| TOS: Irri | 2 | 15.9 | 3.3 | 0.063 |
| Season: Pop | 3 | 214.8 | 2.1 | 0.096 |
| TOS: Pop | 6 | 211.8 | 1.3 | 0.265 |
| Irri: Pop | 3 | 200.6 | 0.3 | 0.834 |
| Season: Genotype | 5 | 210.3 | 4.8 | 0.000^***^ |
| TOS: Genotype | 10 | 205.0 | 2.6 | 0.005^**^ |
| Irri: Genotype | 5 | 194.5 | 0.5 | 0.771 |
| Pop: Genotype | 15 | 196.3 | 1.2 | 0.273 |
| Season: TOS: Irri | 2 | 23.5 | 5.3 | 0.013^*^ |
| Season: TOS: Pop | 6 | 229.4 | 0.9 | 0.476 |
| Season: Irri: Pop | 3 | 239.0 | 2.1 | 0.105 |
| TOS: Irri: Pop | 6 | 195.1 | 2.1 | 0.056 |
| Season: TOS: Genotype | 10 | 224.3 | 3.1 | 0.001^**^ |
| Season: Irri: Genotype | 5 | 213.4 | 1.6 | 0.165 |
| TOS: Irri: Genotype | 10 | 140.7 | 0.9 | 0.565 |
| Season: Pop: Genotype | 15 | 209.3 | 1.4 | 0.130 |
| TOS: Pop: Genotype | 30 | 214.0 | 1.0 | 0.514 |
| Irri: Pop: Genotype | 15 | 204.6 | 1.4 | 0.148 |
| Season: TOS: Irri: Pop | 6 | 233.8 | 2.3 | 0.035^*^ |
| Season: TOS: Irri: Genotype | 10 | 224.5 | 1.3 | 0.216 |
| Season: TOS: Pop: Genotype | 27 | 201.3 | 0.8 | 0.805 |
| Season: Irri: Pop: Genotype | 15 | 196.3 | 0.9 | 0.591 |
| TOS: Irri: Pop: Genotype | 30 | 212.7 | 0.5 | 0.995 |
| Season: TOS: Irri: Pop: Genotype | 12 | 211.2 | 0.8 | 0.658 |
| Where ^*^, ^**^ and ^***^ represents significance at the ≤ 0.05, ≤ 0.01 and ≤ 0.001 levels, respectively. | | | | |

| **Table S4.** Fixed effects testing results from the linear mixed model fitted with residual maximum likelihood (REML) for predicting grain weight for the late winter, spring, and summer sown sorghum in the 2019-2020 and 2020-2021 seasons using season, time of sowing (TOS), irrigation (Irri), plant population density (Pop) and Genotype as fixed factors and replicates and main plots as random effects. | | | | |
| --- | --- | --- | --- | --- |
| Model parameter | Df | denDF | F-Value | *p*-Value |
| (Intercept) | 1 | 3.1 | 1869.0 | 0.000^***^ |
| Season | 1 | 3.2 | 71.9 | 0.003^**^ |
| TOS | 2 | 14.5 | 15.2 | 0.000^***^ |
| Irri | 1 | 14.6 | 0.7 | 0.408 |
| Pop | 3 | 172.4 | 5.8 | 0.000^***^ |
| Genotype | 5 | 170.9 | 23.4 | 0.000^***^ |
| Season: TOS | 2 | 12.0 | 10.4 | 0.002^**^ |
| Season: Irri | 1 | 11.8 | 2.0 | 0.180 |
| TOS: Irri | 2 | 15.0 | 1.5 | 0.265 |
| Season: Pop | 3 | 171.1 | 0.5 | 0.655 |
| TOS: Pop | 6 | 171.7 | 4.5 | 0.000^***^ |
| Irri: Pop | 3 | 174.5 | 4.3 | 0.006^**^ |
| Season: Genotype | 5 | 167.2 | 8.5 | 0.000^***^ |
| TOS: Genotype | 10 | 177.3 | 6.0 | 0.000^***^ |
| Irri: Genotype | 5 | 179.0 | 0.9 | 0.465 |
| Pop: Genotype | 15 | 180.0 | 0.6 | 0.882 |
| Season: TOS: Irri | 2 | 12.7 | 1.0 | 0.397 |
| Season: TOS: Pop | 6 | 186.2 | 2.5 | 0.025^*^ |
| Season: Irri: Pop | 3 | 191.1 | 0.6 | 0.607 |
| TOS: Irri: Pop | 6 | 191.4 | 1.1 | 0.392 |
| Season: TOS: Genotype | 10 | 168.7 | 2.5 | 0.008^**^ |
| Season: Irri: Genotype | 5 | 178.4 | 1.2 | 0.335 |
| TOS: Irri: Genotype | 10 | 193.4 | 0.6 | 0.803 |
| Season: Pop: Genotype | 15 | 182.6 | 1.6 | 0.087 |
| TOS: Pop: Genotype | 30 | 172.0 | 0.7 | 0.836 |
| Irri: Pop: Genotype | 15 | 226.6 | 0.7 | 0.832 |
| Season: TOS: Irri: Pop | 6 | 196.0 | 1.1 | 0.385 |
| Season: TOS: Irri: Genotype | 10 | 185.9 | 0.5 | 0.877 |
| Season: TOS: Pop: Genotype | 30 | 173.1 | 1.0 | 0.498 |
| Season: Irri: Pop: Genotype | 15 | 185.7 | 0.8 | 0.724 |
| TOS: Irri: Pop: Genotype | 30 | 199.5 | 0.6 | 0.933 |
| Season: TOS: Irri: Pop: Genotype | 19 | 184.7 | 1.2 | 0.254 |

Where ^*^, ^**^ and ^***^ represents significance at the ≤ 0.05, ≤ 0.01 and ≤ 0.001 levels, respectively.

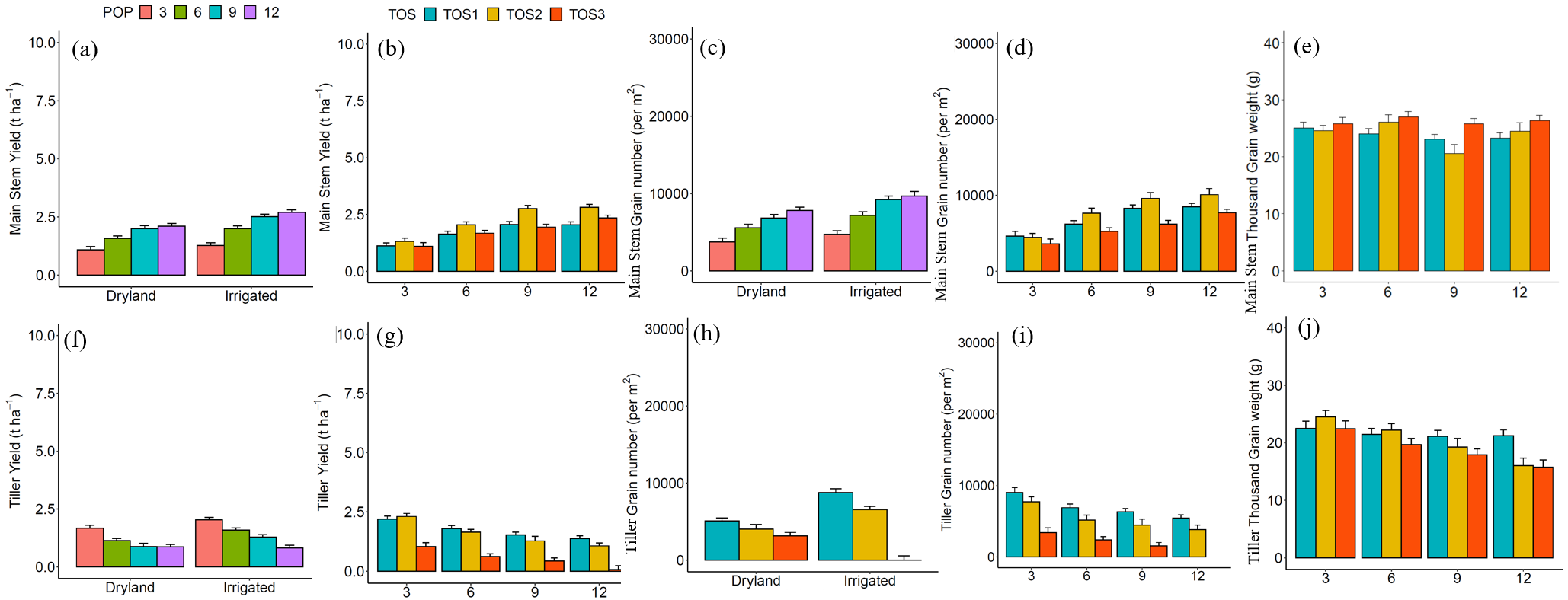

Figure S4. Effects of (a) irrigation (i.e., dryland and irrigation) by plant population density (i.e., 3, 6, 9 and 12 pl m^2^) and (b) the time of sowing (TOS, i.e., late winter-TOS1, spring-TOS2 and summer-TOS3) by plant population density on main stem yield (t ha^-1^); (c) irrigation by plant population density and (d) TOS by plant population density on main stem grain number (per m^2^); (e) TOS by plant population density on main stem thousand-grain weight; (f) irrigation by plant population density and (g) TOS by plant population density on tiller yield (t ha^-1^); (h) irrigation by TOS; (i) TOS by plant population density on tiller grain numbers (per m^2^) and (j) TOS by plant population density on tiller thousand-grain weight across 2019-2020 and 2020-2021 seasons at Nangwee, Queensland, Australia.

| **Table S5.** Fixed effects testing results from the linear mixed model fitted with residual maximum likelihood (REML) for predicting main stem grain yield for the late winter, spring, and summer sown sorghum in the 2019-2020 and 2020-2021 seasons using season, time of sowing (TOS), irrigation (Irri), plant population density (Pop) and Genotype as fixed factors and replicates and main plots as random design effects. | | | | |
| --- | --- | --- | --- | --- |
| Model parameter | Df | denDF | F-Value | *p*-Value |
| (Intercept) | 1 | 3.7 | 702.7 | 0.000^***^ |
| Season | 1 | 3.9 | 169.2 | 0.000^***^ |
| TOS | 2 | 16.5 | 15.9 | 0.000^***^ |
| Irri | 1 | 17.0 | 24.8 | 0.000^***^ |
| Pop | 3 | 223.9 | 89.8 | 0.000^***^ |
| Genotype | 5 | 230.2 | 16.5 | 0.000^***^ |
| Season: TOS | 2 | 17.2 | 2.8 | 0.087 |
| Season: Irri | 1 | 17.3 | 1.2 | 0.290 |
| TOS: Irri | 2 | 17.4 | 0.5 | 0.600 |
| Season: Pop | 3 | 316.5 | 39.2 | 0.000^***^ |
| TOS: Pop | 6 | 240.0 | 3.3 | 0.004^**^ |
| Irri: Pop | 3 | 248.4 | 4.0 | 0.009^**^ |
| Season: Genotype | 5 | 305.4 | 6.0 | 0.000^***^ |
| TOS: Genotype | 10 | 245.3 | 3.8 | 0.000^***^ |
| Irri: Genotype | 5 | 251.0 | 1.2 | 0.319 |
| Pop: Genotype | 15 | 251.2 | 1.4 | 0.150 |
| Season: TOS: Irri | 2 | 17.6 | 0.3 | 0.733 |
| Season: TOS: Pop | 6 | 324.3 | 1.0 | 0.448 |
| Season: Irri: Pop | 3 | 328.1 | 0.3 | 0.823 |
| TOS: Irri: Pop | 6 | 272.9 | 1.4 | 0.231 |
| Season: TOS: Genotype | 10 | 325.1 | 3.9 | 0.000^***^ |
| Season: Irri: Genotype | 5 | 326.1 | 2.1 | 0.062 |
| TOS: Irri: Genotype | 10 | 275.2 | 0.5 | 0.877 |
| Season: Pop: Genotype | 15 | 327.8 | 1.0 | 0.484 |
| TOS: Pop: Genotype | 30 | 265.5 | 0.8 | 0.800 |
| Irri: Pop: Genotype | 15 | 273.0 | 0.9 | 0.529 |
| Season: TOS: Irri: Pop | 6 | 334.5 | 1.2 | 0.310 |
| Season: TOS: Irri: Genotype | 10 | 329.8 | 1.0 | 0.438 |
| Season: TOS: Pop: Genotype | 30 | 322.3 | 1.4 | 0.078 |
| Season: Irri: Pop: Genotype | 15 | 332.1 | 0.9 | 0.597 |
| TOS: Irri: Pop: Genotype | 30 | 295.1 | 0.4 | 0.998 |
| Season: TOS: Irri: Pop: Genotype | 28 | 326.5 | 1.4 | 0.106 |

Where ^*^, ^**^ and ^***^ represents significance at the ≤ 0.05, ≤ 0.01 and ≤ 0.001 levels, respectively.

| **Table S6.** Fixed effects testing results from the linear mixed model fitted with residual maximum likelihood (REML) for predicting main stem grain number for the late winter, spring, and summer sown sorghum in the 2019-2020 and 2020-2021 seasons using season, time of sowing (TOS), irrigation (Irri), plant population density (Pop) and Genotype as fixed factors and replicates and main plots as random design effects. | | | | |
| --- | --- | --- | --- | --- |
| Model parameter | Df | denDF | F-Value | *p*-Value |
| (Intercept) | 1 | 11.3 | 1631.0 | 0.000^***^ |
| Season | 1 | 3.1 | 154.5 | 0.001^**^ |
| TOS | 2 | 17.1 | 15.9 | 0.000^***^ |
| Irri | 1 | 16.3 | 25.1 | 0.000^***^ |
| Pop | 3 | 248.9 | 82.8 | 0.000^***^ |
| Genotype | 5 | 234.2 | 10.2 | 0.000^***^ |
| Season: TOS | 2 | 19.1 | 0.0 | 0.971 |
| Season: Irri | 1 | 18.3 | 0.2 | 0.702 |
| TOS: Irri | 2 | 17.0 | 0.0 | 0.980 |
| Season: Pop | 3 | 289.3 | 21.5 | 0.000^***^ |
| TOS: Pop | 6 | 262.1 | 2.2 | 0.041^*^ |
| Irri: Pop | 3 | 252.7 | 2.2 | 0.084 |
| Season: Genotype | 5 | 294.9 | 7.8 | 0.000^***^ |
| TOS: Genotype | 10 | 248.4 | 4.0 | 0.000^***^ |
| Irri: Genotype | 5 | 232.6 | 0.5 | 0.800 |
| Pop: Genotype | 15 | 250.9 | 1.0 | 0.419 |
| Season: TOS: Irri | 2 | 25.3 | 0.4 | 0.672 |
| Season: TOS: Pop | 6 | 309.0 | 1.0 | 0.459 |
| Season: Irri: Pop | 3 | 307.3 | 0.3 | 0.837 |
| TOS: Irri: Pop | 6 | 282.8 | 1.9 | 0.086 |
| Season: TOS: Genotype | 10 | 298.0 | 3.0 | 0.001^**^ |
| Season: Irri: Genotype | 5 | 303.2 | 1.6 | 0.169 |
| TOS: Irri: Genotype | 10 | 274.3 | 0.8 | 0.607 |
| Season: Pop: Genotype | 15 | 296.9 | 1.0 | 0.486 |
| TOS: Pop: Genotype | 30 | 265.3 | 0.9 | 0.560 |
| Irri: Pop: Genotype | 15 | 127.3 | 1.2 | 0.298 |
| Season: TOS: Irri: Pop | 6 | 309.8 | 0.3 | 0.920 |
| Season: TOS: Irri: Genotype | 10 | 303.0 | 0.5 | 0.902 |
| Season: TOS: Pop: Genotype | 28 | 292.7 | 0.7 | 0.882 |
| Season: Irri: Pop: Genotype | 15 | 298.1 | 0.7 | 0.824 |
| TOS: Irri: Pop: Genotype | 30 | 282.7 | 0.4 | 0.999 |
| Season: TOS: Irri: Pop: Genotype | 22 | 298.3 | 1.0 | 0.454 |

Where ^*^, ^**^ and ^***^ represents significance at the ≤ 0.05, ≤ 0.01 and ≤ 0.001 levels, respectively.

| **Table S7.** Fixed effects testing results from the linear mixed model fitted with residual maximum likelihood (REML) for predicting main stem grain weight for the late winter, spring, and summer sown sorghum in the 2019-2020 and 2020-2021 seasons using season, time of sowing (TOS), irrigation (Irri), plant population density (Pop) and Genotype as fixed factors and replicates and main plots as random design effects. | | | | |
| --- | --- | --- | --- | --- |
| Model parameter | Df | denDF | F-Value | *p*-Value |
| (Intercept) | 1 | 2.9 | 2683.0 | 0.000^***^ |
| Season | 1 | 2.8 | 70.4 | 0.004^**^ |
| TOS | 2 | 14.6 | 24.4 | 0.000^***^ |
| Irri | 1 | 14.1 | 0.0 | 0.953 |
| Pop | 3 | 225.1 | 11.6 | 0.000^***^ |
| Genotype | 5 | 225.1 | 38.5 | 0.000^***^ |
| Season: TOS | 2 | 9.4 | 13.3 | 0.002^**^ |
| Season: Irri | 1 | 9.5 | 0.5 | 0.504 |
| TOS: Irri | 2 | 14.2 | 2.1 | 0.156 |
| Season: Pop | 3 | 128.1 | 1.4 | 0.233 |
| TOS: Pop | 6 | 219.3 | 4.2 | 0.000^***^ |
| Irri: Pop | 3 | 228.4 | 2.4 | 0.072 |
| Season: Genotype | 5 | 126.4 | 9.3 | 0.000^***^ |
| TOS: Genotype | 10 | 217.6 | 7.2 | 0.000^***^ |
| Irri: Genotype | 5 | 221.0 | 2.5 | 0.033^*^ |
| Pop: Genotype | 15 | 222.3 | 1.2 | 0.276 |
| Season: TOS: Irri | 2 | 10.3 | 0.5 | 0.610 |
| Season: TOS: Pop | 6 | 133.2 | 1.3 | 0.248 |
| Season: Irri: Pop | 3 | 145.6 | 0.3 | 0.841 |
| TOS: Irri: Pop | 6 | 222.5 | 1.0 | 0.397 |
| Season: TOS: Genotype | 10 | 150.5 | 1.8 | 0.070 |
| Season: Irri: Genotype | 5 | 146.7 | 1.2 | 0.330 |
| TOS: Irri: Genotype | 10 | 218.7 | 0.5 | 0.875 |
| Season: Pop: Genotype | 15 | 140.0 | 1.1 | 0.338 |
| TOS: Pop: Genotype | 30 | 209.0 | 1.1 | 0.404 |
| Irri: Pop: Genotype | 15 | 209.6 | 0.3 | 0.992 |
| Season: TOS: Irri: Pop | 6 | 151.4 | 1.2 | 0.328 |
| Season: TOS: Irri: Genotype | 10 | 159.9 | 0.6 | 0.795 |
| Season: TOS: Pop: Genotype | 30 | 147.6 | 0.8 | 0.701 |
| Season: Irri: Pop: Genotype | 15 | 157.3 | 0.6 | 0.896 |
| TOS: Irri: Pop: Genotype | 30 | 199.5 | 1.1 | 0.333 |
| Season: TOS: Irri: Pop: Genotype | 22 | 156.8 | 0.9 | 0.598 |

Where ^*^, ^**^ and ^***^ represents significance at the ≤ 0.05, ≤ 0.01 and ≤ 0.001 levels, respectively.

| **Table S8.** Fixed effects testing results from the linear mixed model fitted with residual maximum likelihood (REML) for predicting tiller grain yield for the late winter-, spring-, and summer- sown sorghum in the 2019-2020 and 2020-2021 seasons using season, time of sowing (TOS), irrigation (Irri), plant population density (Pop) and Genotype as fixed factors and replicates and main plots as random design effect. | | | | |
| --- | --- | --- | --- | --- |
| Model parameter | Df | denDF | F-Value | *p*-Value |
| (Intercept) | 1 | 3.0 | 206.2 | 0.001^**^ |
| Season | 1 | 3.0 | 315.5 | 0.000^***^ |
| TOS | 2 | 58.5 | 37.3 | 0.000^***^ |
| Irri | 1 | 53.1 | 22.7 | 0.000^***^ |
| Pop | 3 | 244.0 | 27.6 | 0.000^***^ |
| Genotype | 5 | 243.6 | 7.9 | 0.000^***^ |
| Season: TOS | 2 | 10.5 | 27.3 | 0.000^***^ |
| Season: Irri | 1 | 10.4 | 10.2 | 0.009^**^ |
| TOS: Irri | 2 | 52.6 | 2.2 | 0.117 |
| Season: Pop | 3 | 277.8 | 16.0 | 0.000^***^ |
| TOS: Pop | 6 | 247.0 | 5.5 | 0.000^***^ |
| Irri: Pop | 3 | 249.0 | 0.9 | 0.429 |
| Season: Genotype | 5 | 270.3 | 8.6 | 0.000^***^ |
| TOS: Genotype | 10 | 241.7 | 0.9 | 0.562 |
| Irri: Genotype | 5 | 246.1 | 0.5 | 0.809 |
| Pop: Genotype | 15 | 247.7 | 1.2 | 0.296 |
| Season: TOS: Irri | 2 | 10.8 | 21.3 | 0.000^***^ |
| Season: TOS: Pop | 6 | 279.9 | 1.4 | 0.210 |
| Season: Irri: Pop | 3 | 284.2 | 2.5 | 0.063 |
| TOS: Irri: Pop | 6 | 253.7 | 0.5 | 0.815 |
| Season: TOS: Genotype | 10 | 273.2 | 3.2 | 0.001^**^ |
| Season: Irri: Genotype | 5 | 274.6 | 1.7 | 0.127 |
| TOS: Irri: Genotype | 10 | 250.9 | 0.5 | 0.854 |
| Season: Pop: Genotype | 15 | 262.1 | 1.0 | 0.448 |
| TOS: Pop: Genotype | 30 | 244.1 | 0.8 | 0.708 |
| Irri: Pop: Genotype | 15 | 245.7 | 0.5 | 0.954 |
| Season: TOS: Irri: Pop | 6 | 275.8 | 1.9 | 0.088 |
| Season: TOS: Irri: Genotype | 10 | 270.8 | 1.5 | 0.123 |
| Season: TOS: Pop: Genotype | 30 | 256.1 | 0.7 | 0.915 |
| Season: Irri: Pop: Genotype | 15 | 263.1 | 1.7 | 0.046^*^ |
| TOS: Irri: Pop: Genotype | 30 | 275.6 | 0.5 | 0.991 |
| Season: TOS: Irri: Pop: Genotype | 25 | 261.2 | 1.3 | 0.130 |

Where ^*^, ^**^ and ^***^ represents significance at the ≤ 0.05, ≤ 0.01 and ≤ 0.001 levels, respectively.

| **Table S9.** Fixed effects testing results from the linear mixed model fitted with residual maximum likelihood (REML) for predicting tiller grain number for the late winter-, spring-, and summer- sown sorghum in the 2019-2020 and 2020-2021 seasons using season, time of sowing (TOS), irrigation (Irri), plant population density (Pop) and Genotype as fixed factors and replicates and main plots as random design effects. | | | | |
| --- | --- | --- | --- | --- |
| Model parameter | Df | denDF | F-Value | *p*-Value |
| (Intercept) | 1 | 11.1 | 911.7 | 0.000^***^ |
| Season | 1 | 3.5 | 195.8 | 0.000^***^ |
| TOS | 2 | 11.2 | 48.7 | 0.000^***^ |
| Irri | 1 | 10.1 | 25.7 | 0.000^***^ |
| Pop | 3 | 197.7 | 22.9 | 0.000^***^ |
| Genotype | 5 | 205.8 | 7.8 | 0.000^***^ |
| Season: TOS | 2 | 15.9 | 22.4 | 0.000^***^ |
| Season: Irri | 1 | 16.0 | 4.0 | 0.062 |
| TOS: Irri | 2 | 11.3 | 8.8 | 0.005^**^ |
| Season: Pop | 3 | 254.4 | 5.1 | 0.002^**^ |
| TOS: Pop | 6 | 216.6 | 3.7 | 0.002^**^ |
| Irri: Pop | 3 | 214.0 | 0.8 | 0.518 |
| Season: Genotype | 5 | 247.3 | 4.4 | 0.001^**^ |
| TOS: Genotype | 10 | 216.2 | 1.0 | 0.465 |
| Irri: Genotype | 5 | 192.5 | 0.1 | 0.990 |
| Pop: Genotype | 15 | 216.1 | 1.4 | 0.134 |
| Season: TOS: Irri | 2 | 24.8 | 19.5 | 0.000^***^ |
| Season: TOS: Pop | 6 | 269.0 | 0.8 | 0.568 |
| Season: Irri: Pop | 3 | 278.0 | 1.6 | 0.198 |
| TOS: Irri: Pop | 6 | 236.4 | 0.6 | 0.741 |
| Season: TOS: Genotype | 10 | 263.4 | 2.9 | 0.002^**^ |
| Season: Irri: Genotype | 5 | 269.7 | 0.9 | 0.490 |
| TOS: Irri: Genotype | 10 | 232.3 | 0.7 | 0.726 |
| Season: Pop: Genotype | 15 | 258.0 | 1.3 | 0.237 |
| TOS: Pop: Genotype | 30 | 220.8 | 1.0 | 0.422 |
| Irri: Pop: Genotype | 15 | 133.9 | 1.1 | 0.410 |
| Season: TOS: Irri: Pop | 6 | 271.6 | 2.0 | 0.063 |
| Season: TOS: Irri: Genotype | 10 | 264.2 | 1.0 | 0.463 |
| Season: TOS: Pop: Genotype | 30 | 249.0 | 0.7 | 0.889 |
| Season: Irri: Pop: Genotype | 15 | 250.5 | 0.9 | 0.569 |
| TOS: Irri: Pop: Genotype | 30 | 259.1 | 0.4 | 0.998 |
| Season: TOS: Irri: Pop: Genotype | 20 | 250.0 | 1.0 | 0.512 |

Where ^*^, ^**^ and ^***^ represents significance at the ≤ 0.05, ≤ 0.01 and ≤ 0.001 levels, respectively.

| **Table S10.** Fixed effects testing results from the linear mixed model fitted with residual maximum likelihood (REML) for predicting tiller grain weight for the late winter-, spring-, and summer- sown sorghum in the 2019-2020 and 2020-2021 seasons using season, time of sowing (TOS), irrigation (Irri), plant population density (Pop) and Genotype as fixed factors and replicates and main plots as random design effects. | | | | |
| --- | --- | --- | --- | --- |
| Model parameter | Df | denDF | F-Value | *p*-Value |
| (Intercept) | 1 | 3.7 | 1119.0 | 0.000^***^ |
| Season | 1 | 3.6 | 134.6 | 0.001^**^ |
| TOS | 2 | 16.6 | 0.7 | 0.493 |
| Irri | 1 | 16.7 | 0.1 | 0.817 |
| Pop | 3 | 245.5 | 11.6 | 0.000^***^ |
| Genotype | 5 | 246.6 | 9.7 | 0.000^***^ |
| Season: TOS | 2 | 15.7 | 20.0 | 0.000^***^ |
| Season: Irri | 1 | 15.9 | 0.5 | 0.502 |
| TOS: Irri | 2 | 16.9 | 0.5 | 0.637 |
| Season: Pop | 3 | 191.7 | 7.5 | 0.000^***^ |
| TOS: Pop | 6 | 249.3 | 3.1 | 0.006^**^ |
| Irri: Pop | 3 | 249.0 | 2.0 | 0.119 |
| Season: Genotype | 5 | 185.1 | 15.0 | 0.000^***^ |
| TOS: Genotype | 10 | 252.0 | 4.6 | 0.000^***^ |
| Irri: Genotype | 5 | 254.0 | 0.3 | 0.926 |
| Pop: Genotype | 15 | 252.3 | 1.2 | 0.297 |
| Season: TOS: Irri | 2 | 16.8 | 1.8 | 0.190 |
| Season: TOS: Pop | 6 | 198.2 | 7.7 | 0.000^***^ |
| Season: Irri: Pop | 3 | 208.1 | 1.8 | 0.142 |
| TOS: Irri: Pop | 6 | 254.6 | 0.2 | 0.976 |
| Season: TOS: Genotype | 10 | 210.5 | 8.7 | 0.000^***^ |
| Season: Irri: Genotype | 5 | 207.9 | 0.6 | 0.736 |
| TOS: Irri: Genotype | 10 | 255.7 | 1.0 | 0.414 |
| Season: Pop: Genotype | 15 | 198.6 | 1.2 | 0.307 |
| TOS: Pop: Genotype | 30 | 243.9 | 0.9 | 0.630 |
| Irri: Pop: Genotype | 15 | 245.9 | 1.4 | 0.130 |
| Season: TOS: Irri: Pop | 6 | 208.7 | 1.4 | 0.220 |
| Season: TOS: Irri: Genotype | 10 | 214.9 | 2.0 | 0.037 |
| Season: TOS: Pop: Genotype | 30 | 200.8 | 1.4 | 0.091 |
| Season: Irri: Pop: Genotype | 15 | 215.9 | 1.0 | 0.453 |
| TOS: Irri: Pop: Genotype | 30 | 249.9 | 0.9 | 0.584 |
| Season: TOS: Irri: Pop: Genotype | 23 | 211.1 | 1.2 | 0.247 |

Where ^*^, ^**^ and ^***^ represents significance at the ≤ 0.05, ≤ 0.01 and ≤ 0.001 levels, respectively.
